# Pervasive Somatic Mutations in *NF1* Drive Cell Type and Spatially Specific Clonal Selection in Neurofibromatosis

**DOI:** 10.64898/2026.09.29.755447

**Authors:** Daniel A. Snellings, Kow Essuman, Benjamin Finander, Eitan Goodman, Christie Cambridge, Jannis Buecking, Brian Chhouk, Chunhui Cai, Liang Sun, Urmi Ghosh, Michael B. Miller, Christopher A. Walsh

## Abstract

Tumorigenesis is a multi-step process shaped by the accumulation of somatic driver mutations, but identifying the earliest steps and cell of origin of tumorigenesis remains challenging. Single cell genomics in >1.7 million single cells across donors with the cancer predisposition syndrome neurofibromatosis type 1 (NF-1), shows that somatic ‘second-hit’ mutations in *NF1* are pervasive and show positive clonal selection in oligodendrocytes (OLs) and OL precursor cells (OPCs), but are exceedingly rare in neurons and other lineages. Combined spatial genotyping and transcriptomic analysis enabled dissection of cell autonomous and non-autonomous effects of second-hit mutations including clonal expansion of mutant OPCs with strong bias for cerebral gray matter versus white matter, suggesting that gray matter influences OPC proliferation and differentiation via non-cell autonomous interactions between OPCs and neurons. *NF1*-null OL lineage cells upregulate *CDKN2A*—a tumor suppressor frequently lost in *NF1*-mutant tumors— whereas brain cancers from these same individuals show *CDKN2A* loss across multiple cancer cell states, suggesting *CDKN2A* loss in OPCs is a critical step in tumor progression. Our findings reveal developmentally early, exquisitely lineage-specific *NF1* somatic mutations in premalignant NF-1 brain, suggest cell autonomous and non-autonomous roles in clonal selection in brain, and suggest that clones harboring cancer driver mutations can persist over a lifetime in phenotypically normal tissues.

## Introduction

Somatic mutations are central to cancer initiation and progression, and in recent years, the discovery of cancer driver mutations in normal human tissue has redefined our understanding of the origins of tumorigenesis ^1–5^. However, discerning the functional impact of these early mutations in normal human tissue remains a challenge. Individuals with tumor predisposition syndromes caused by germline mutation in a tumor suppressor gene provide an opportunity to study the effects of somatic mutations in phenotypically normal human tissue. In these individuals, the germline mutation alone is insufficient for transformation, but primes the cells for tumorigenesis following subsequent somatic driver mutations that arise throughout life.

Neurofibromatosis type 1 (NF-1) is the most common human brain tumor predisposition syndrome, where germline loss of the *NF1* gene leads to tumors in the central and peripheral nervous system, as well as varied non-tumor phenotypes in the brain and other organ systems ^6,7^. *NF1* is also commonly mutated in non-syndromic gliomas ^8–10^, and no curative therapies exist either for NF-1, or for gliomas ^6,11^. A recent report identified somatic loss of *NF1* in histologically normal tissues from multiple individuals with NF-1^12^. Though this study only assayed brain tissue in one donor, it suggests that biallelic mutation of *NF1* may be common in the brain of individuals with NF-1. Here, we sought to track the frequency and lineage specificity of these second hits to understand how both germline and somatic mutation of *NF1* affect mutant clonal dynamics, gene expression, and chromatin accessibility, that ultimately may lead to gliomagenesis.

## Results

### Widespread mosaic loss of NF1 in the brain

Somatic mutations resulting in biallelic loss of *NF1* were identified in fresh-frozen post-mortem samples of 7 non-tumor regions across the brain from 10 individuals with NF-1 and 8 healthy controls (Fig. 1A). Independent genetic testing of NF-1 donors confirmed the presence of a deleterious germline mutation in *NF1* (table S1). Analysis of these NF-1 brains by a neuropathologist also revealed no histologically detectable tumor (fig. S1).

**Figure 1:**
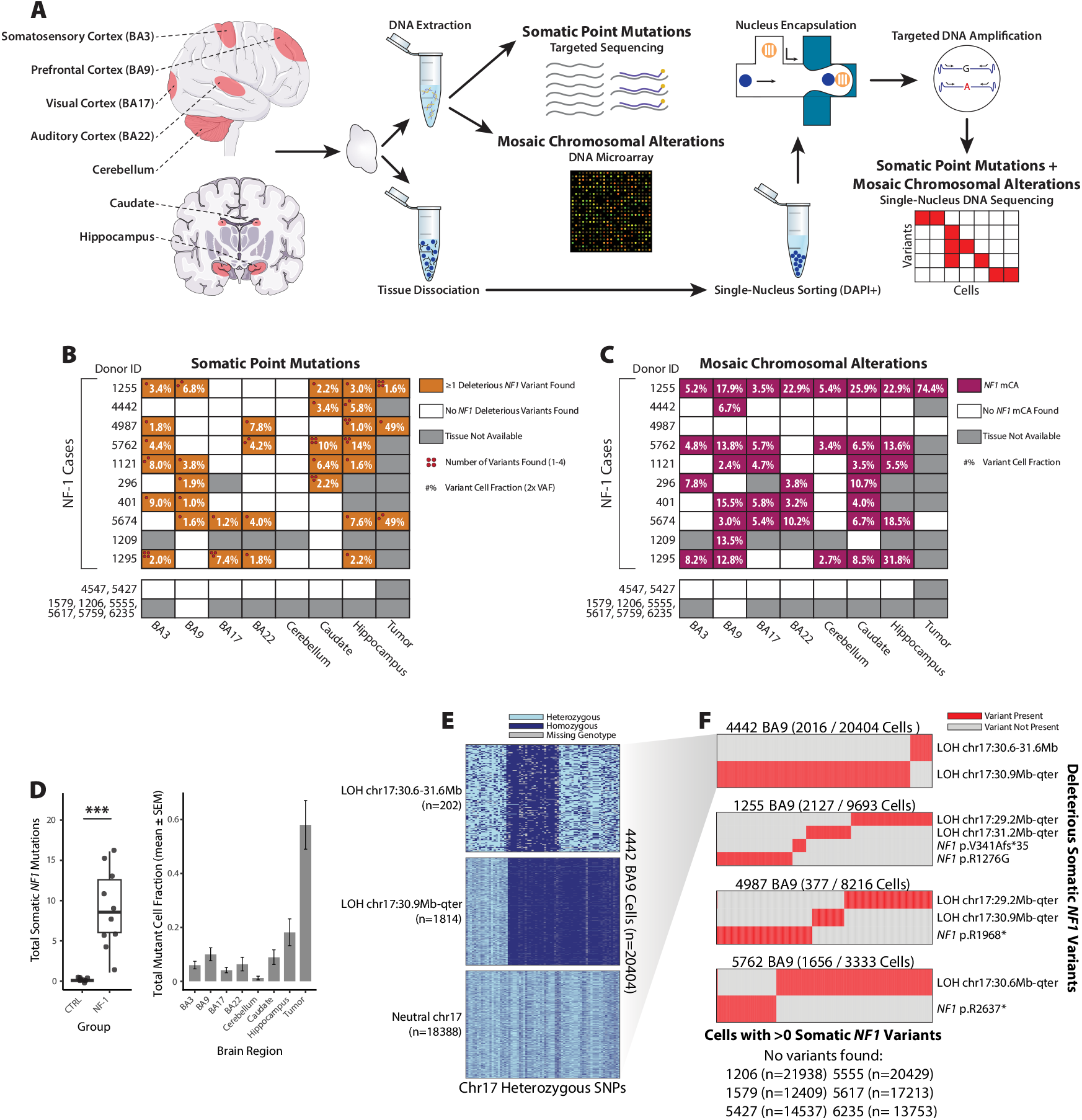
Multifocal *NF1*-null clones in the brain of humans with neurofibromatosis type 1. (**A**) Experimental diagram. (**B**) Heatmap of nonsynonymous *NF1* somatic point mutations identified in all samples. The inset percentage corresponds to the summed cell fractions (allele frequency * 2) for all somatic variants identified in the sample. The number of variants identified in each sample is indicated by the number of red pips in the top left of each box. No variants were identified in samples from neurotypical controls, therefore multiple individuals are grouped per row. (**C**) Heatmap of mCA events impacting chr17. (**D**) Enrichment of total mutations (point mutations and mCA combined) in individuals with NF-1 compared to controls (*** p < 0.0005, two-tailed Wilcoxon test), and enrichment of somatic mutations in NF-1 individuals by brain region. (**E**) Cell x variant heatmap from scDNA-seq data from individual 4442. The color of each tile in the heatmap indicates the zygosity of the variant where homozygous variants may be either homozygous for the reference or alternate allele. (**F**) Clonal structure of somatic variants causing *NF1* LOF. The total number of mutated cells and total number of cells assayed is noted for each sample. No deleterious *NF1* mutations were identified in 6 neurotypical controls.

Targeted duplex sequencing of a panel of 201 genes including *NF1* (table S2), to a mean depth of 21,594x (416x duplex depth) (fig. S2A-B), identified multiple deleterious somatic point mutations in *NF1* in non-tumor tissue from 9/10 NF-1 donors. Single tissue samples often contained multiple deleterious point mutations (Fig. 1B, table S3), with none identified in control samples. The allele frequency of somatic *NF1* point mutations ranged from 0.3% to 5.1% (mean 1.7% ±1.1%, table S3). Notably, five samples from three unrelated individuals harbored identical 1bp deletions in an exonic homopolymer resulting in a frameshift (fig. S2C). Closer evaluation of this site revealed duplex reads supporting this variant were present in nearly all NF-1 samples, but not controls (fig. S2D), suggesting that indels in this homopolymer are a common and recurrent mechanism of somatic *NF1* inactivation.

Large-scale mosaic chromosomal alterations (mCAs) disrupting *NF1*, identified using MoChA analysis ^13,14^ of microarray data for each sample, revealed even more frequent inactivation of *NF1* by loss of heterozygosity (LOH). MoChA identified mCAs affecting *NF1* in 9/10 NF-1 donors, frequently present across multiple brain regions (Fig. 1C, table S4), but absent in controls. Interestingly, mCAs in non-tumor NF-1 brain tissue were detected at significantly greater cell fractions than point mutations (mean 9.6% ±7.4% vs. mean 3.4% for point mutations, *p* = 5.2 × 10^−7^; linear mixed-effects model). MoChA identified most events as copy-neutral LOH. Several gains and losses were identified, though most were present at low cell fractions which is known to decrease the reliability of copy number determination with MoChA ^15^ (table S4). Combining both point mutations and mCA, *NF1* loss is pervasive in all 10 individuals with NF-1 across multiple brain regions (Fig. 1D).

### Somatic NF1 mutations are polyclonal

Amplicon-based single-cell DNA sequencing using the Tapestri system (Fig. 1A) revealed that in tissue samples with multiple somatic *NF1* mutations, the mutations are strictly mutually exclusive within cells, showing that the NF-1 brain is comprised of a heterogeneous mixture of many independent *NF1*-null clones. To detect both point mutations and LOH in single cells, we developed a panel of primers to amplify the exons of *NF1* in addition to 262 common polymorphic sites across chromosome 17 (table S5). This panel was applied to nuclei from prefrontal cortex samples of 4 individuals with NF-1 and 6 controls assaying a total of 142,057 nuclei. LOH was detected by germline heterozygous variants shifting to homozygosity across a contiguous genomic region (Fig. 1E). This strategy achieved greater sensitivity than microarray analysis, identifying LOH down to 0.7% of cells and revealing that 3 of the 4 NF-1 samples harbored multiple LOH events with distinct breakpoints undetected by bulk microarray analysis, all resulting in loss of the functional *NF1* allele. In addition, 4 deleterious somatic point mutations in *NF1* were identified across 3 samples (Fig. 1F). No LOH or deleterious *NF1* point mutations were called in the 6 control samples, though 4/6 controls showed possible low-level LOH in <0.3% of cells (fig. S3), possibly reflecting artifactual allelic dropout or the baseline burden of LOH in healthy individuals. In each sample with multiple somatic events disrupting *NF1*, the events were mutually exclusive between cells, with only 9/41,646 exceptions, likely reflecting cell doublets (Fig. 1F). These results demonstrate that *NF1* loss occurred multiple times in independent lineages, giving rise to a polyclonal tissue comprised of a mixture of cells with either heterozygous or homozygous loss of *NF1*.

### Cell type-specific somatic loss of NF1

Analysis of the cell types carrying somatic mutations disrupting *NF1* showed that they were overwhelmingly in the oligodendroglial lineage. Single-nucleus RNA-sequencing (snRNA-seq) on samples of prefrontal cortex from ten individuals with NF-1 and four control donors enabled genotyping of *NF1* LOH in each cell using Numbat ^16^, a computational tool that leverages variant phasing information to detect LOH using germline heterozygous variants (Fig. 2A). Numbat determined chromosome 17 LOH status (neutral or LOH, posterior probability >0.99) in >60% of 215,025 nuclei encompassing the expected distribution of cell types (Fig. 2B-C, fig S4A-B). The vast majority (>95%) of LOH calls were in OLs and OPCs (Fig. 2D), and these were the only cell types with significant (FDR < 0.05) enrichment of LOH relative to controls (Fig. 2E). Analysis of 75,782 additional nuclei sorted with the oligodendroglial lineage marker Sox10 confirmed the high burden of LOH in OLs and OPCs (Fig. 2F, fig. S4C-E). Microglia showed a non-significant increase in LOH (FDR = 0.094) in NF-1 samples driven by a high burden of LOH in a subset of samples. While functional effects of heterozygous *NF1* mutation have been previously documented in microglia ^17,18^, LOH in microglia suggests that homozygous loss of *NF1* may contribute to microglia-dependent phenotypes in NF-1. LOH was not detected in neurons, which was confirmed by an additional 85,732 nuclei sorted with the neuronal lineage marker NeuN (Fig. 2G, fig. S4F-H). Overall, these data indicate a strong bias for LOH in the oligodendroglial lineage, suggesting either a lineage-specific vulnerability to LOH events, or lineage-specific positive selection for *NF1* loss.

**Figure 2:**
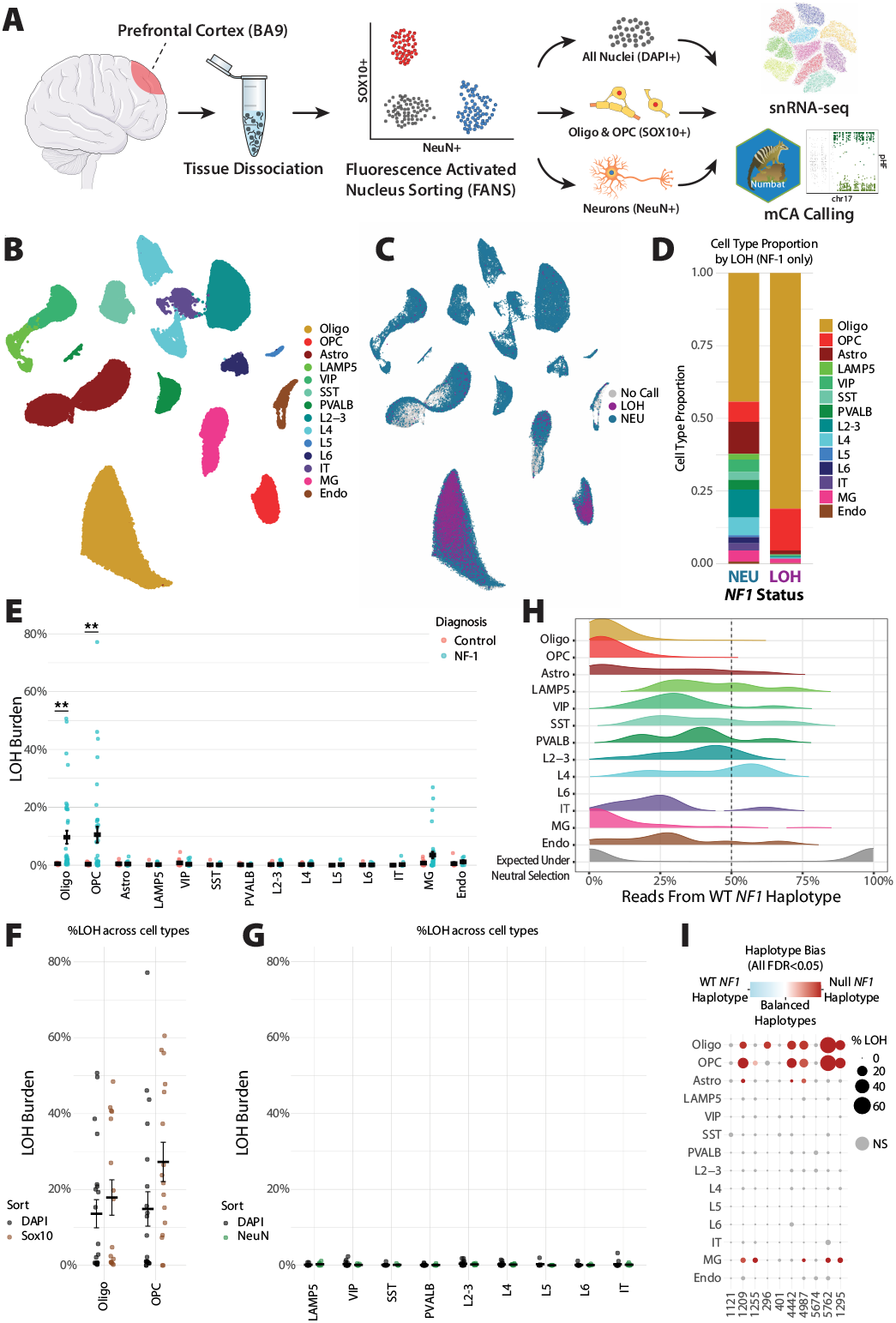
LOH is strongly enriched in the oligodendroglial lineage. (**A**) Experimental diagram. (**B**) UMAP representation of cell types present in BA9 snRNA-seq samples. (**C**) UMAP representation of cells colored by LOH status (Blue = neutral, Purple = LOH, Gray = no call). (**E**) % Chr17q LOH burden for all cell types, NF-1 and neurotypical samples compared. P-values are BH-adjusted, two-sided t-test values. (** p < 0.005). (**F-G**) Comparison of LOH burdens in Sox10 (F) and NeuN (G) sorted and non-sorted snRNA-seq samples across cell types. (**H**) Distribution of haplotype balance in cells identified to have LOH by Numbat for each cell types from NF-1 donors. Haplotypes are oriented such that 0% corresponds to all reads being derived from the *NF1-*null haplotype, while 100% corresponds to all reads being derived from the WT *NF1* haplotype. (**I**) Haplotype bias split across donor and cell type. Colored dots denote significantly biased haplotype distributions (FDR < 0.05.

Somatic mutations in *NF1* showed strong clonal selection in the oligodendrocyte and microglial lineages, but not the neuronal lineage. The ratio of synonymous to nonsynonymous mutations (dN/dS) measures selection ^19^ on single-nucleotide variants, but similar metrics are not well defined for LOH, therefore we developed an analogous measure to detect selection on LOH. Under neutral selection, LOH would impact both haplotypes equally leading to equal numbers of cells homozygous for the *NF1*-null allele and the functional *NF1* allele, as observed in snDNA-seq experiments from control donors (~0.3% per allele, fig. S3). Quantification of the allelic imbalance in each cell with LOH revealed near-complete bias for the *NF1-*null allele in OLs, OPCs, and microglia as well as significant bias (p < 0.05, BH-adjusted binomial test) in astrocytes from a subset of donors (Fig. 2H-I, fig. S4I). In contrast, the infrequent LOH events called in neurons lacked haplotype bias indicating these events are largely false positives. These data are consistent with observations from mouse models of NF-1^20^ and show that biallelic loss of *NF1* is under positive selection in OLs, OPCs, microglia, and astrocytes, but not in neurons.

### Clonal expansion of NF1-null OPCs in NF-1 cortex

Spatial analysis revealed OPC expansion in the prefrontal cortex of NF-1 donors consistent with previous studies in mice ^20,21^. Multiplex error robust fluorescent in situ hybridization (MERFISH) ^22^ of 5 sections of non-tumor prefrontal cortex from 3 control donors and 8 sections from 5 NF-1 donors (Fig. 3A), using a custom 140-gene panel (table S6) targeting major cell types in the human brain, enabled gray/white matter segmentation (Fig. 3B, fig. S5A). Analysis of 800,634 cells focused on the gray matter revealed a two-fold increase in the proportion of OPCs in NF-1 versus controls (FDR<0.05) (Fig. 3C). Interestingly, astrocytes were also expanded in NF-1 compared to controls, despite very low levels of astrocyte LOH in snRNA-seq data (Fig. 2E), implying non-cell autonomous effects. There was no significant change in neurons.

**Figure 3:**
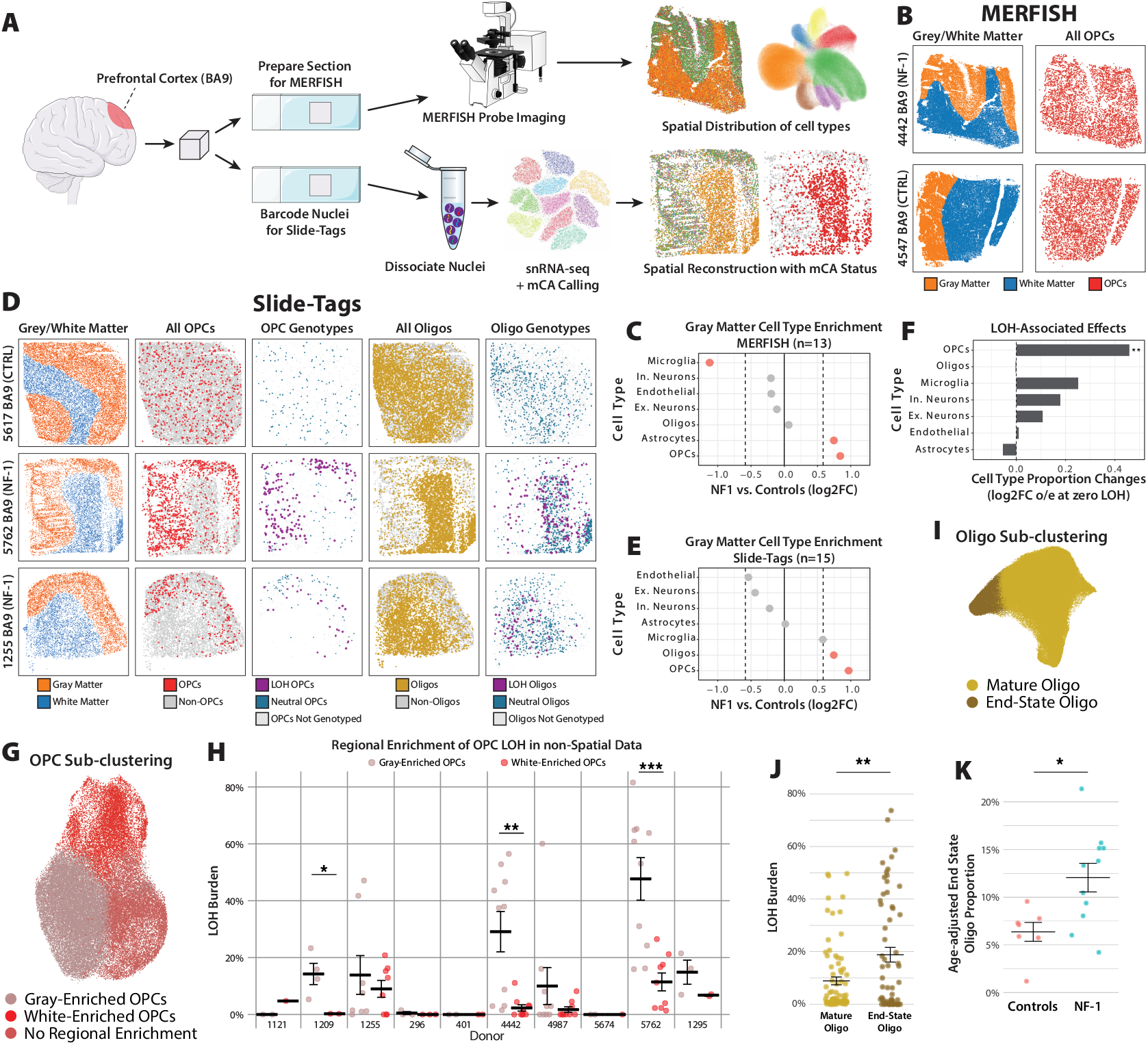
Expansion of the oligodendroglial lineage in NF-1 cortex. (**A**) Experimental diagram. (**B**) Representative spatial reconstructions from MERFISH data of NF1 and control samples. (**C**) Cell type enrichment and depletion in MERFISH data from NF-1 vs controls in gray matter from all 13 sections. (**D**) Representative images from 2 NF-1 donors (high and low levels of LOH) and 1 control donor. Panels show gray/white matter classifications and the spatial distribution of OPCs and OLs as well as LOH genotypesas determined by Numbat. (**E**) Cell type enrichment and depletion in Slide-tags data from NF-1 vs controls in gray matter from all 15 sections. (**F**) Shift in gray matter cell type proportions associated with the burden of LOH (** p < 0.005, linear model: proportion ~ %LOH * region). (**G**) UMAP representation of all OPCs across all datasets clustered independently. **(H)** % Chr17q LOH burden for OPC subtypes in non-spatial data, split by individual donor (* p < 0.05, ** p < 0.005, p < 0.0005). (**I**) UMAP representation of all OLs across all datasets clustered independently. (**J**) % Chr17q LOH burden for OL subtypes in all data. (**K**) Age-adjusted end-state OL proportions in control and NF1 brain samples.

Spatial transcriptomics with paired *in situ* genotyping of *NF1* LOH using Slide-tags ^23^ revealed striking localization of mutant OPCs in gray matter compared to white matter. Slide-tags on sections of prefrontal cortex from 10 NF-1 donors, and 5 control donors (each section textsubscript 1 cm^2^) yielded 315,888 high-quality cells (fig. S5B-C). As in previous experiments, Numbat detected LOH events disrupting *NF1* with single-cell resolution, enabling tracking of spatial distributions of mutated cells (Fig. 3D). Slide-tags confirmed the two-fold increase (FDR<0.05) in OPCs in the gray matter of NF-1 donors versus controls (Fig. 3E). The enrichment of other cell types differs between methods, potentially reflecting heterogeneity in NF-1 as observed in the snRNA-seq experiments (Fig. 2I). The overall burden of LOH was similar between gray and white matter (p = 0.47); however, the cell type composition differed greatly, with striking enrichment of LOH OPCs in gray matter versus white matter (OR=8.05, p<0.0001, Fig. 3D). To estimate the impact of LOH on cell type composition in the gray matter, we fit donor-level LOH and cell type proportion data to linear models for each cell type and compared the expected proportions at the zero-LOH intercept to the observed proportions. This analysis revealed that LOH is associated with a} 1.4-fold increase of OPCs in the gray matter (log_2_FC=0.48, p=0.0008) accounting for 43% of the observed increase between NF-1 donors and controls (Fig. 3F). The remaining 57% may be attributed to point mutations in *NF1* that cannot be detected with this method or an effect of *NF1* heterozygosity, independent of somatic mutations.

While the increase in LOH OPCs in gray matter was driven by one donor (35% in OPCs from donor 5762 vs. mean of 2.7% in other NF-1 donors), analysis of a larger sample set by using spatial data to identify distinct transcriptional signatures of gray vs white matter OPCs (fig. S5D-E) in non-spatial data (Fig. 3G, fig. S5F-G) confirmed increased gray matter OPCs in NF-1 compared to controls and a strong bias for LOH in gray matter OPCs across multiple donors (Fig. 3H, fig. S5H). Whereas the spatial data identified distinct DEGs in gray matter OPCs and white matter OPCs consistent with reduced differentiation signaling and increased neuronal signaling in gray matter (fig. S5D-E), sub-clustering OPCs in the non-spatial data revealed populations of OPCs marked by expression of these regionally biased DEGs (Fig. 3G, fig. S5F-G), confirmed as gray/white-enriched by co-clustering with spatial data (fig. S5H). Pooled data shows increased gray-enriched OPCs in NF-1 compared to controls with >90% of LOH OPCs in the gray-enriched subcluster, consistent across multiple donors (p = 3.03e-5) (Fig. 3H, fig. S5H). The increased OPCs in NF-1 and the robust, consistent bias for mutant OPCs in the gray matter may reflect non-cell autonomous impacts on OPC proliferation and differentiation in response to neuronal activity, as has been observed in mouse models of NF-1^24^.

Consistent with defective OPC proliferation or differentiation in NF-1, end-state OLs are enriched in NF-1 versus controls and further enriched in LOH OLs versus neutral OLs. Subclustering of OLs across all datasets identified a mature end-state OL cluster marked by decreased *OPALIN* and increased *RBFOX1* expression, consistent with a previous suggestion that these cells may be more mature with reduced capacity for myelination ^25^ (Fig. 3I, fig. S5I). End-state OLs demonstrated increased LOH burden (Fig. 3J), suggesting failure of OPCs with LOH to produce sufficient new OLs. Consistent with impaired differentiation of OPCs, the age-adjusted proportion of end-state OLs is significantly higher in NF-1 than in controls (Fig. 3K), indicative of reduced OL turnover overall in NF-1. The increase of end-state OLs in NF-1 may contribute to myelin abnormalities common in humans with NF-1^26^ and myelin decompaction observed in mouse models of NF-1^27^.

### Cell type-specific effects of somatic NF1 mutations on gene expression

To analyze cell type-specific effects of somatic *NF1* mutations, we focused initially on donors without brain tumors to avoid potential confounding effects on gene expression from the cancer state suggested by principal component analysis (fig. S6A). Excluding samples from donors diagnosed with tumors markedly decreased the number of differentially expressed genes (DEGs, FDR < 0.05 and —log_2_FC— > 0.5) (fig. S6B-E). While the etiology underlying this difference in DEGs between the two NF-1 groups remains to be determined, samples with brain tumors may have received chemotherapy and/or radiation treatment, and resultant gene expression changes may reflect additional shared effects of dying cells associated with brain tumors.

OPCs and OLs from NF-1 samples demonstrate gene expression patterns indicative of impaired differentiation. Mature OLs from NF-1 samples show gene expression signatures consistent with increased end-state OL proportion in NF-1, including dysregulated expression of cadherins ^25^ *CDH1* and *CDH13, OPALIN*, and *BMP8A* (Fig. 4A, data S1).

**Figure 4:**
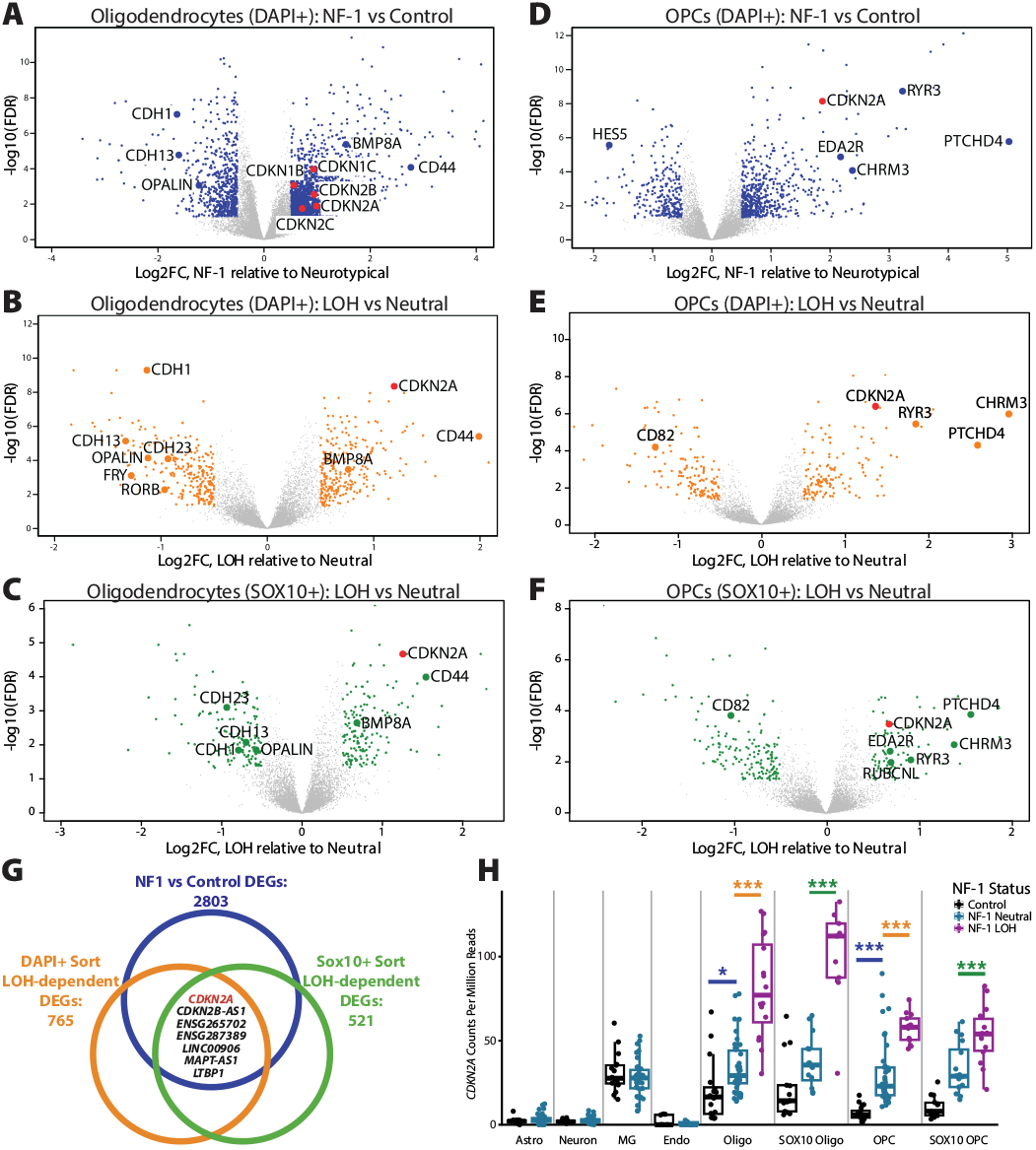
*CDKN2A* and OL differentiation gene dysregulation after both germline and somatic *NF1* loss in OLs and OPCs. (**A-F**) Volcano plots for OLs and OPCs comparing samples from individuals with NF-1 to neurotypical samples (panels A, D) and cells from NF-1 donors with LOH compared to cells without LOH (Neutral) (panels B, C, E, F). *CDKN2A* and significant OL differentiation genes highlighted. (**G**) Venn diagram of DGE results demonstrating that *CDKN2A* is one of 7 differentially expressed genes (| log_2_(FC|) > 0.5 and FDR < 0.05) in OLs and OPCs comparing NF-1 to neurotypical samples, and LOH NF-1 nuclei to neutral NF-1 nuclei for both the unsorted snRNA-seq dataset and the SOX10-sorted dataset. (**H**) EdgeR CPM (Counts Per Million mapped reads) expression values of *CDKN2A* across cell types pseudo-bulked by sample across snRNA-seq datasets demonstrate both significantly increased expression with germline loss of *NF1* and significantly increased expression with somatic loss of *NF1*. Significance determined using edgeR pseudo-bulk differential gene expression with BH multiple hypothesis correction, threshold for significance was |log_2_(FC)| > 0.5 and FDR < 0.05.

Additionally, several senescence genes were increased in NF-1 OLs, including *CDKN1B, CDKN1C, CDKN2A, CDKN2B, CDKN2C*, and *CD44*. Several of these genes are further increased in OLs with LOH, indicating that *NF1* dosage impacts gene expression (Fig. 4B-C, fig. S7G, data S2). Similarly, NF-1 OPCs show increased expression of immature OPC genes ^28^ *RYR3, CHRM3*, and *PTCHD4* and senescence genes like *CDKN2A* and *EDA2R* that are often further increased in OPCs with LOH (Fig. 4D-F, fig. S7H). Overall, the DEGs from NF-1 donors vs. control donors, and LOH cells vs. neutral cells both highlight impaired differentiation and senescence in the oligodendroglial lineage.

Cell-cell communication analysis using CellChat ^29^ highlighted multiple differential ligand-receptor interactions consistent with a coordinated shift toward myelin repair that was broadly distributed across many cell types in NF-1 brain, independent of LOH status (fig. S7A-D). Of 18 significant ligand-receptor interactions, 9 have established roles in myelin dynamics: ANGPTL ^30^, SPP1^31^, VISTA^32^, GAS^33^, CX3C^34^, PSAP ^35^, APP ^36^, SEMA6^37^, and NT ^38^. Signaling from astrocytes and microglia appear to be central players in the differential cell-cell communication network in NF-1 (fig. S7B-D). Astrocytes and microglia also have increased expression of immune reactivity genes ^39–41^ *CD81, SOCS3, TLR2*, and *TMEM163* (fig. S7E-F, data S1), overlapping with known functions in myelin clearance ^42^. Notably, these changes are predominantly in cell types with low levels of *NF1* LOH, suggesting these effects are likely a secondary response to dysfunction in the oligodendroglial lineage, rather than a cell-autonomous effect of *NF1* loss.

### Increased CDKN2A expression in heterozygous and NF1-null cells

Overlapping mutation-dependent DEGs in OLs and OPCs across multiple datasets converged on a dosage-dependent upregulation of the tumor suppressor *CDKN2A*, which is a potential compensatory mechanism that inhibits tumorigenesis in *NF1*-null cells. Only 7 genes are consistently significant when comparing NF-1 to control and LOH to neutral OLs and OPCs (Fig. 4G). This set of 7 genes includes the tumor suppressor and senescence marker gene *CDKN2A*, the deletion of which is common in high grade gliomas ^43^, and is an essential genetic event in the transformation of malignant peripheral nerve sheath tumors ^44,45^, which are common in NF-1^46,47^. *CDKN2A* expression in OPCs and OLs significantly increases as *NF1* dosage decreases (Fig. 4H, data S1-2). *CDKN2A* is a key negative feedback mechanism in the *RAS/MAPK* pathway ^48^ previously shown to be increased in human fibroblasts from NF-1 donors ^49^. Taken together, increased expression of *CDKN2A* after both germline and somatic loss of *NF1* could explain how OPCs compensate for lost tumor suppression function, succumb to senescence, and fail to differentiate normally in NF-1, allowing glial expansion but suppressing tumorigenesis, and making subsequent loss of *CDKN2A* strongly tumorigenic.

### CDKN2A loss is associated with more aggressive cell states in NF-1 glioblastoma

Single-nucleus multiome sequencing (RNA + ATAC) of paired tumor and normal brain tissue from NF-1 donors revealed that loss of *CDKN2A* is specific to *NF1*-null clones with malignant features. Out of 10 NF-1 donors, 5 were diagnosed with brain tumors, and 3 had available highquality tumor-infiltrated tissue. Donors 1255 and 4987 were diagnosed with glioblastoma multiforme (GBM) and 5674 with anaplastic astrocytoma. Multiome data from 71,244 nuclei (Fig. 5A-B, fig. S8A-B) identified large cell clusters (designated as ‘presumed malignant’) with canonical malignant features including high expression of *CD44* and *VIM*, increased accessibility of *FOS* and *JUN* motifs (fig. S8C-D), and multiple CNVs including homozygous loss of *CDKN2A* which were highly specific to tumor-infiltrated samples (fig. S9A-B). Tumors from donor 4987 and 5674 harbored deleterious point mutations in *NF1* rather than *NF1* LOH (Fig. 1B, table S3), preventing detection using Numbat, but in the tumor from donor 1255, Numbat identified identical *NF1* LOH events shared by both non-tumor oligodendrocytes and presumed malignant cells, consistent with the hypothesis that the tumor cell of origin was an *NF1*-null cell in the oligodendrogilal lineage (Fig. 5C, fig. S9A). Lineage reconstruction in this sample with Numbat identified that *CDKN2A* loss was exclusive to presumed malignant, *NF1*-null cells (fig. S9A), and in all tumors, cells with *CDKN2A* loss were only present as a subset of presumed malignant cells. Presumed malignant cells with *CDKN2A* loss have more aggressive transcriptional states than cells with *CDKN2A* intact. *CDKN2A* contained several of the top linked peaks (peaks of accessible chromatin with strong positive correlations with gene expression levels across cells) ^50^ (fig. S9C), making *CDKN2A* a strong candidate to influence malignant cell states previously characterized in glioblastoma and astrocytoma ^51–53^. *NF1-*mutated tumors have previously been linked to mesenchymal-like (MES-like) states in glioblastoma ^8,54^, with the MES2-like state in particular associated with more aggressive tumor phenotypes ^51^. Strikingly, the GBM samples exhibit up to a seven-fold increase in the proportion of MES2-like states in cells with *CDKN2A* loss (Fig. 5F-G, fig. S8E-G). The anaplastic astrocytoma was enriched in both astrocyte-like (AC-like) and OPC-like cell states previously observed in astrocytoma ^52^ (fig. S8F) and also demonstrated *CDKN2A*-dependent clustering and cell state (Fig. 5F-G). Malignant cells with *CDKN2A* loss in this anaplastic astrocytoma were much more likely to have OPC-like cell states than AC-like cell states (Fig. 5F). Overall, *CDKN2A* loss only occurs in *NF1*-null clones, and is correlated with more aggressive malignant cell states in NF-1-associated tumors.

**Figure 5:**
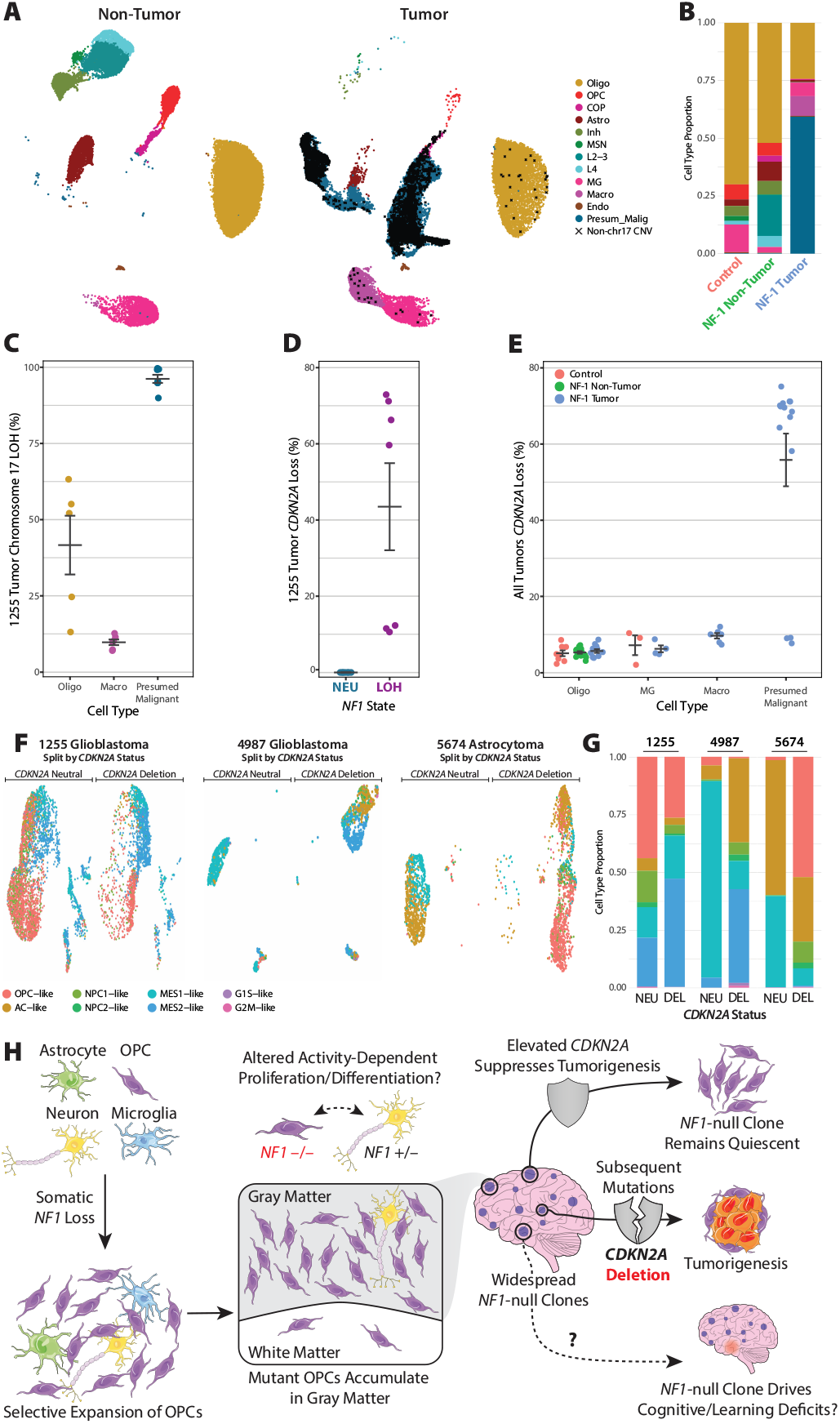
*CDKN2A* loss drives malignant cells towards more aggressive cell states in NF-1-associated tumors. (**A**) UMAP representation of cell types present in Multiome dataset, split by whether sample is a tumor, cells with non-chromosome 17 CNVs labeled with a black X. (**B**) Cell types present in neurotypical, NF-1 BA9, and NF-1 tumor samples. (**C**) % Chr17q LOH burden for selected cell types in sample 1255 GBM. (**D**) % *CDKN2A* loss burden in presumed malignant nuclei in 1255 GBM, split by chromosome 17 status. (**E**) % *CDKN2A* loss burden in select cell types, split by sample type. (**F**) Subclustering of presumed malignant nuclei in each of the 3 tumors in the dataset, split by *CDKN2A* status. (**G**) Malignant cell states in each tumor, split by *CDKN2A* status. (**H**) Model of the role of *NF1* somatic mutations in the pathogenesis of NF-1. At the cell-level, loss of *NF1* drives selective clonal expansion of OPCs. Mutant OPCs are strongly biased the gray matter, likely due to interactions with neurons leading to altered activity-dependent proliferation and/or differentiation. The majority of NF1-null clones in the brain remain quiescent due to mutation-dependent upregulation of *CDKN2A* expression, while a small subset of clones acquire additional mutations, including CDKN2A deletion, that drive tumorigenesis. We speculate that *NF1*-null clones may also contribute to cognitive phenotypes in NF-1, even though they do not progress to bona-fide tumors.

## Discussion

Our integrative analysis of somatic mutations in the NF-1 brain uncovered pervasive *NF1*-null clones largely restricted to the oligodendroglial lineage and identifies cell autonomous and non-autonomous effects of *NF1* loss across cell types (Fig. 5H). Our focus on NF-1 provides a unique opportunity to explore how tumorigenic mutations arise, persist, and influence the brain long before neoplastic transformation. Collectively, our results provide a stepwise model of tumor progression and highlight an unexplored axis of NF-1 pathogenesis that may bridge the gap between animal models and the human phenotype.

The pattern of *NF1*-null clones we observe is the result of numerous independent somatic mutations throughout the brain that confer a cell type-dependent selective advantage. Our results show that single brain regions often harbor multiple distinct clones with independent somatic mutations. These clones are dominated by OLs and OPCs with sporadic contributions from astrocytes and microglia driven by positive selection, consistent with observations in mice following deletion of *Nf1* in radial glia ^20^. *NF1*-null clones in the neuronal lineage are rare or absent in the donors studied here, but likely contribute to specific neurological phenotypes like epilepsy in other cases ^55,56^. The striking asymmetry of mutations across cell types may reflect the relative capacity for each cell type to proliferate throughout life ^57^ but may also be influenced by negative selection for *NF1* loss, which our study is not powered to detect. Outside of the brain, second hits have also been observed in café-au-lait spots ^58^, bone ^59,60^, and several neuroectodermal tissues ^12^. Further studies in these tissues may reveal similarly asymmetric clonal selection and identify the essential cell populations driving NF-1-associated phenotypes across the body.

Combining spatial, transcriptional, and mutational information in single cells allowed us to detect both cell autonomous and non-cell autonomous consequences of *NF1* loss. The loss of *NF1* drives cell autonomous clonal expansion in the OL lineage in parallel with seemingly sporadic non-cell autonomous expansion of astrocytes, reminiscent of astrogliosis in NF-1 mice ^61–63^. The strong bias of *NF1*-null OPCs in the gray matter suggests a non-cell autonomous effect from proximity to neurons. We hypothesize that this may be a result of either 1) increased proliferation of mutant OPCs in response to neuronal activity as seen in gliomas ^64,65^; or 2) reduced sensitivity of mutant OPCs to activity-dependent OPC differentiation which has been previously described in NF-1 mice ^24^. Supporting the latter hypothesis, we find that *NF1* loss in OPCs drives cell autonomous transcriptional dysregulation of core genes for OPC differentiation. The loss of *NF1* in OLs and OPCs may also contribute to increased myelin thickness ^27^, hypermyelination, and myelin decompaction ^66^ observed in mouse models of NF-1. Further studies will be required to confirm this link, although the non-cell autonomous transcriptional changes observed in astrocytes and microglia appear consistent with established roles in myelin repair.

Our results point to a stepwise progression of tumor formation starting with a single driver mutation, and suggest that mutant clones may silently persist for long periods in normal tissue. Analysis of GBMs revealed identical somatic mutations shared by malignant cells and non-malignant OLs indicating a direct lineage relationship and suggesting that the abundant clones of *NF1*-null OPCs provide the cell of origin for GBM, consistent with experiments in mouse models ^21^. By extension, the abundance of *NF1*-null clones may represent a quantifiable link to lifetime tumor risk in NF-1. Given that *NF1*-null clones far outnumber brain tumors in our cohort, it is likely that only a small minority of mutant OPCs ever progress to a tumor. Our data suggest that the benign behavior of mutant OPCs may result from compensatory expression of the tumor suppressor *CDKN2A*, which exhibits a dosage-dependent upregulation in *NF1*-null OLs and OPCs. Homozygous deletion of *CDKN2A* is a common event in human cancers ^67^ and is recognized as a key event in the malignant transformation of plexiform neurofibroma in NF-1^44,68^. Consistently, we find that *CDKN2A* loss is associated with more aggressive cell states in NF-1-associated tumors. CDK inhibitors, which mimic the function of *CDKN2A*, are being tested in clinical trials for brain tumors ^69–71^ and are effective in *NF1-*mutant breast cancer ^72^. These drugs may also be effective for the treatment of NF-1-associated brain tumors. Notably, the presence of latent tumorigenic clones in non-tumor brain tissue is not a phenomenon unique to NF-1. Large studies of brain tissue from control donors have identified premalignant clonal selection in ~ 25% of the population ^73,74^. NF-1 offers an amplified window into this process, making it a powerful system to understand how mutation and selection shape the human brain and influence disease risk.

Lastly, second-hit mutations in *NF1* are a compelling candidate to explain the variable expressivity observed in families with NF-1. While heterozygous loss of *NF1* exhibits complete penetrance for NF-1 diagnosis, the severity of the disease varies widely within affected families, including monozygotic twins ^75,76^. Studies of genotype-phenotype correlation have identified few germline variants that drive distinct phenotypes ^77^, suggesting non-inherited factors—such as somatic second-hit mutations—may be a major source of phenotypic variation. Consistent with this hypothesis, studies in mice have found that targeted disruption of *Nf1* in the oligodendroglial lineage leads to motor skill learning deficits ^78–80^, raising the possibility that second-hit mutations may contribute to neurological manifestations that are common, but variable, in individuals with NF-1 (Fig. 5H). The timing and location of second-hit mutations in the oligodendroglial lineage, as well as less common second-hit mutations in other lineages (e.g., neurons or extra-CNS lineages) may be critical factors driving the variable expressivity of NF1-associated phenotypes; however, further studies with deep clinical phenotyping and larger sample sizes will be required to fully understand the impact of second-hit mutations.

## Acknowledgements

We thank Ron Mathieu from the Boston Children’s Hospital Flow Cytometry Core, Shuqiang Li and J. Thomas Janes III for their contribution in scDNA-seq work from the Translational Immunogenomics Lab (TIGL) at Dana-Farber Cancer Institute, Harry Cramer from the Boston Children’s Cellular Imaging Core; RRID:SCR 026485, funded by NIH P50 HD105351, the BIDMC Confocal/Immunostaining Core for assistance with histology, and R.S Hill and J. E. Neil for their administrative help. We also thank Dr. David Miller and Dr. Nicole Ullrich from Boston Children’s Hospital for helpful advice throughout this study. We thank the donors of postmortem tissues for their invaluable contributions to the advancement of science as well as the NIH NeuroBioBank and the Children’s Tumor Foundation for providing tissue samples. D.A.S was supported by F32 NS133029. K.E. was supported by a Hanna Gray Fellowship from the Howard Hughes Medical Institute. M.B.M. is funded by DP2 AG086138 and R01 AG082346. This work was supported by The Dorothy, Emma & Spiro Latsis Benevolent Trust. C.A.W. is also funded by NINDS grants R01NS032457 and UH3NS132138. C.A.W. is an Investigator of the Howard Hughes Medical Institute.

## Author Contributions

D.A.S., K.E., B.F., and C.A.W. conceived of the study. D.A.S. and E.G., performed long-read DNA sequencing and microarray analysis with additional assistance from B.C. for targeted sequencing. D.A.S. and J.B. performed scDNA-seq with Tapestri with additional assistance from C.C. and L.S. for analysis. K.E., E.G., and J.B. performed single-nuclei sorting, snRNA-seq, and multiome experiments. B.F. performed snRNA-seq and multiome analysis with additional assistance from D.A.S. and J.B. for LOH calling with Numbat. K.E., E.G., and C.C., performed MERFISH experiments with additional assistance from C.C. and L.S. for analysis and U.G. for preliminary experiments. D.A.S., C.C., and U.G. performed slide-tags experiments and D.A.S. and B.F. performed the analysis. M.B.M. performed histological analysis from brain tissues. D.A.S., K.E., and B.F. wrote the initial draft of the manuscript. C.A.W. supervised the study and edited the manuscript.

## Swials

### Supplementary Tables

**Table S1. (Supplemental Tables.xlsx)**

Type Cohort demographics and germline *NF1* mutation status.

**Table S2. (Supplemental Tables.xlsx)**

Targeted sites in bulk targeted DNA sequencing panel.

**Table S3. (Supplemental Tables.xlsx)**

Somatic *NF1* point mutations identified by bulk targeted DNA sequencing.

**Table S4. (Supplemental Tables.xlsx)**

Mosaic chromosomal alterations identified by bulk DNA microarray.

**Table S5. (Supplemental Tables.xlsx)**

Targeted sites in snDNA-seq panel.

**Table S6. (Supplemental Tables.xlsx)**

Target genes in MERFISH panel.

### Supplementary Data

**Data S1. (Data S1 NFvsCTRL DEG.xlsx)**

Differentially expressed genes in snRNA-seq comparing NF-1 donors to control donors.

**Data S2. (Data S2 LOHvsNEU DEG.xlsx)**

Differentially expressed genes in snRNA-seq comparing cells with LOH to cells without LOH.

### Supplementary Figures

**Figure S1:**
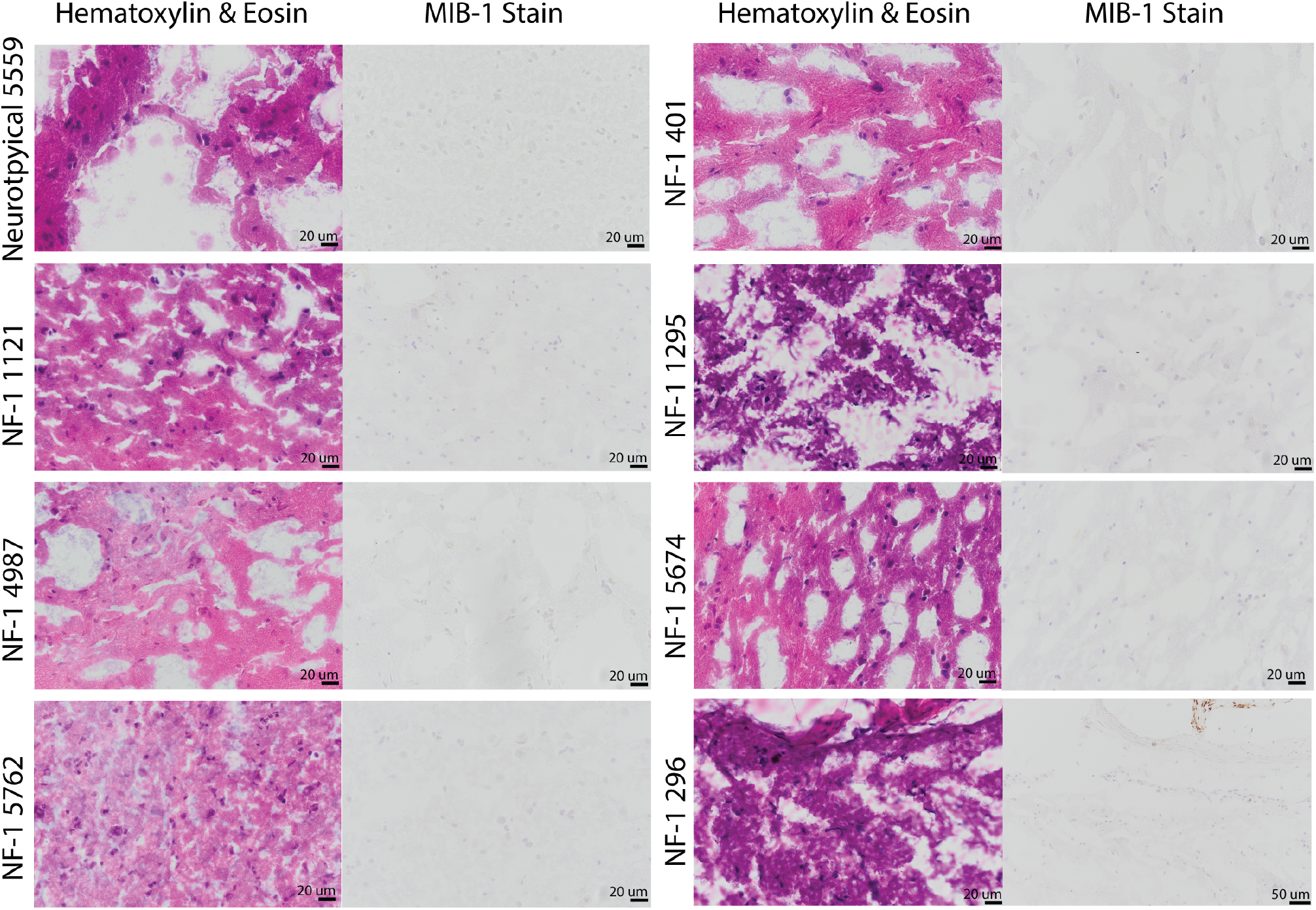
Histopathological analysis of NF-1 brain sections. Hematoxylin and eosin (H&E) and Ki-67/MIB-1 (proliferation marker) stains do not show evidence of hypercellularity or increased proliferation characteristic of brain tumors. Histology was performed on available frozen brain tissue.

**Figure S2:**
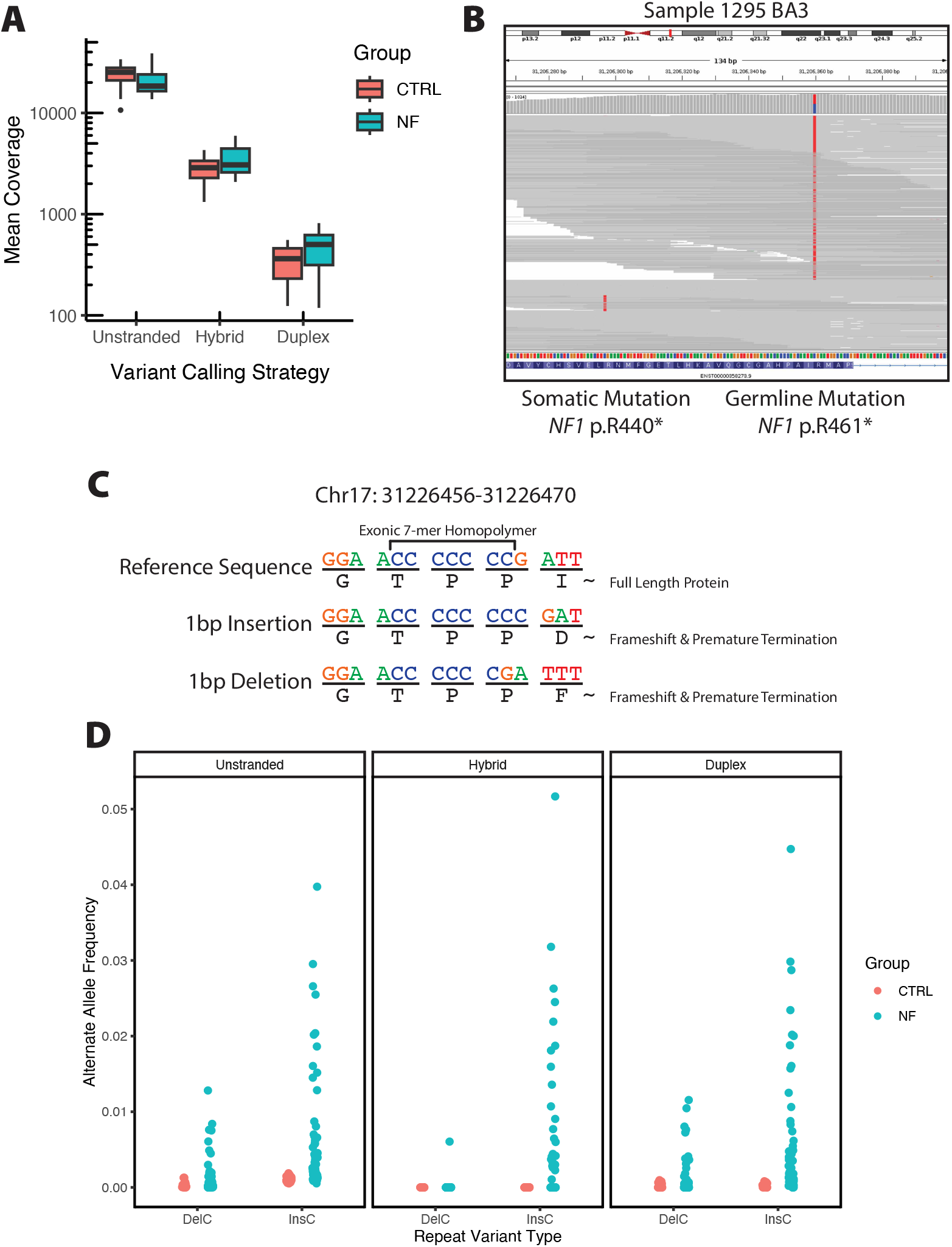
Targeted *NF1* sequencing metrics and presence of recurrent homopolymer mutations. (**A**) Mean coverage in targeted regions for each calling strategy: ignoring strand information (unstranded), using information from both molecules with duplex consensus or consensus from a single strand (hybrid), or strictly considering molecules with duplex consensus (duplex). (**B**) Screenshot from IGV showing a somatic and germline mutation identified in BA3 from 1295. Reads spanning both positions show that the somatic and germline variants are in trans orientation. (**C**) Schematic showing a 7-mer homopolymer that is recurrently mutated in samples from individuals with NF-1. (**D**) Allele frequency for single bp insertions or deletions in the 7-mer homopolymer across calling strategies and disease status.

**Figure S3:**
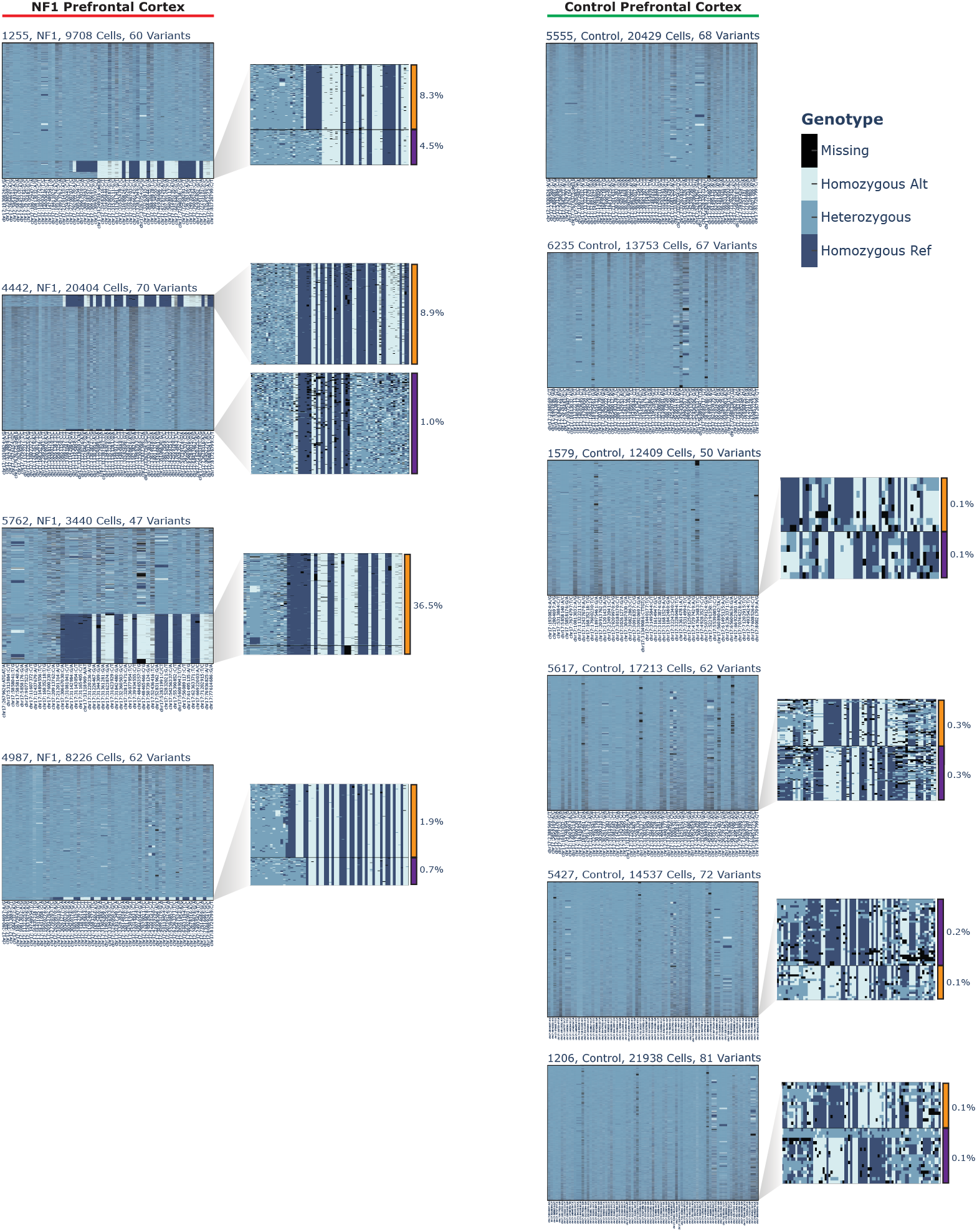
LOH identified in snDNA-seq. Cell (row) x variant (column) matrices are shown for 10 prefrontal cortex (BA9) samples (4 NF-1, 6 control) used for snDNA-seq. Each x axis shows germline heterozygous polymorphic markers on chromosome 17 ordered by genomic coordinate. Each row is a single cell assayed. The fill of each tile in the matrix is the called genotype of that cell; either heterozygous (blue), homozygous for the reference allele (dark blue), homozygous for the alternate allele (light blue), or no genotype was called (black). Eight of the samples also have insets showing a zoomed region of a portion of the parent matrix that show either called LOH events (for NF-1 samples) or possible low level LOH that could not be confidently called (in controls). Note that in the NF-1 samples, the LOH events always impact the same allele (visible by pattern of light/dark blue stripes), whereas the possible LOH in controls is present on both alleles in roughly equal frequency (noted by the inverted stripe pattern between the orange and purple cell populations.

**Figure S4:**
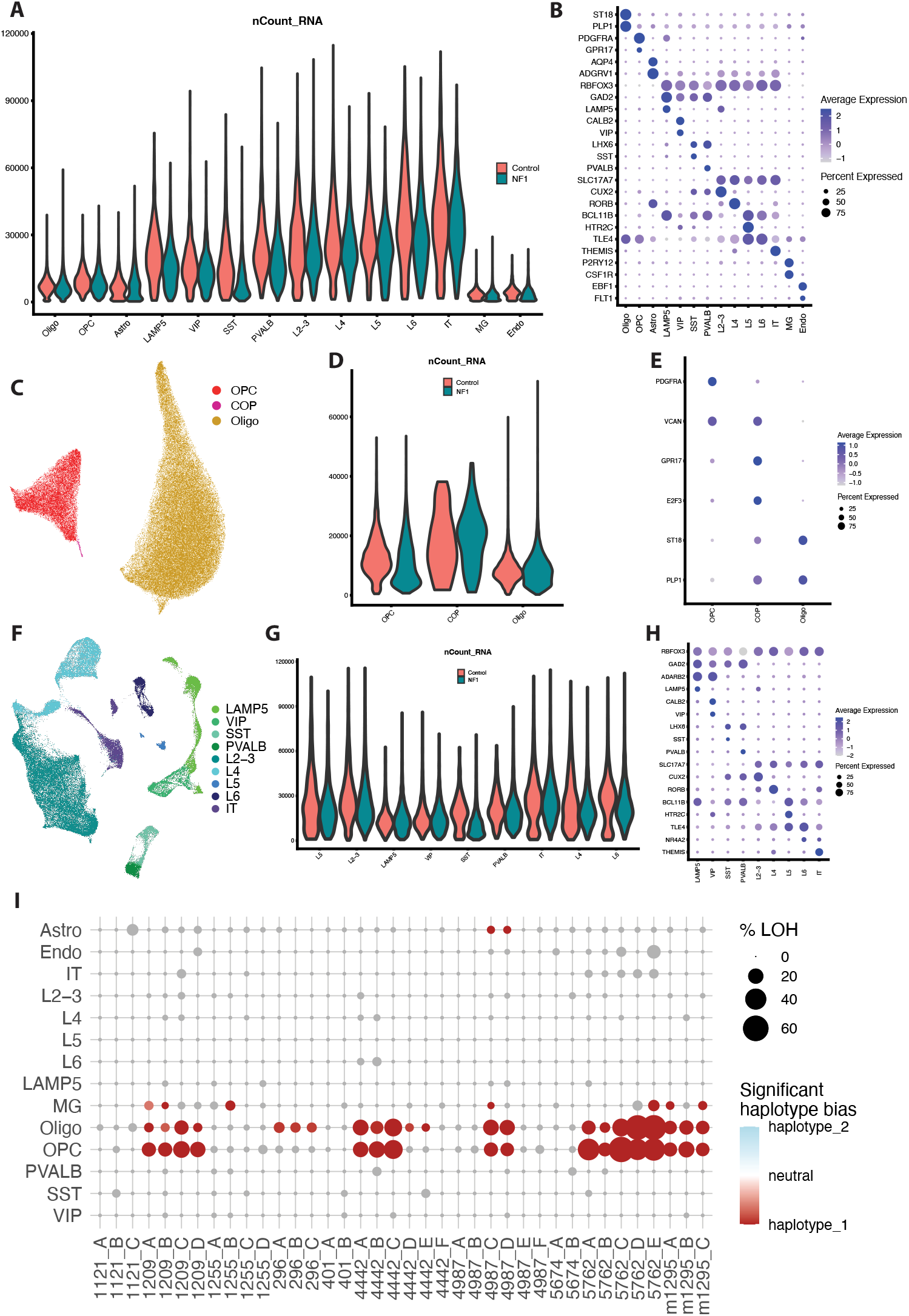
High-quality snRNA-seq data shows sample-level LOH haplotype bias. (**A**) Violin plot of UMI count across cell types in unsorted snRNA-seq dataset, split by NF-1 diagnosis. (**B**) Dot plot showing marker gene expression in all cell types in unsorted snRNA-seq dataset. (**C**) UMAP representation of cell types present in SOX10-sorted experiment. Additional Cell Type Abbreviation: COP = Committed Oligodendrocyte Precursors. (**D**) Violin plot of UMI count across cell types in SOX10 sorted snRNA-seq dataset, split by NF-1 diagnosis. (**E**) Dot plot showing marker gene expression in all cell types in SOX10 sorted snRNA-seq dataset. (**F**) UMAP representation of cell types present in NeuN-sorted experiment. (**G**) Violin plot of UMI count across cell types in NeuN sorted snRNA-seq dataset, split by NF-1 diagnosis. (**H**) Dot plot showing marker gene expression in all cell types in NeuN sorted snRNA-seq dataset. (**I**) Haplotype bias for cells called as harboring LOH split by replicates and cell types. Colored dots denote replicates and cell types with significant haplotype bias (FDR < 0.05, binomial test with Benjamini-Hochberg FDR adjustment).

**Figure S5:**
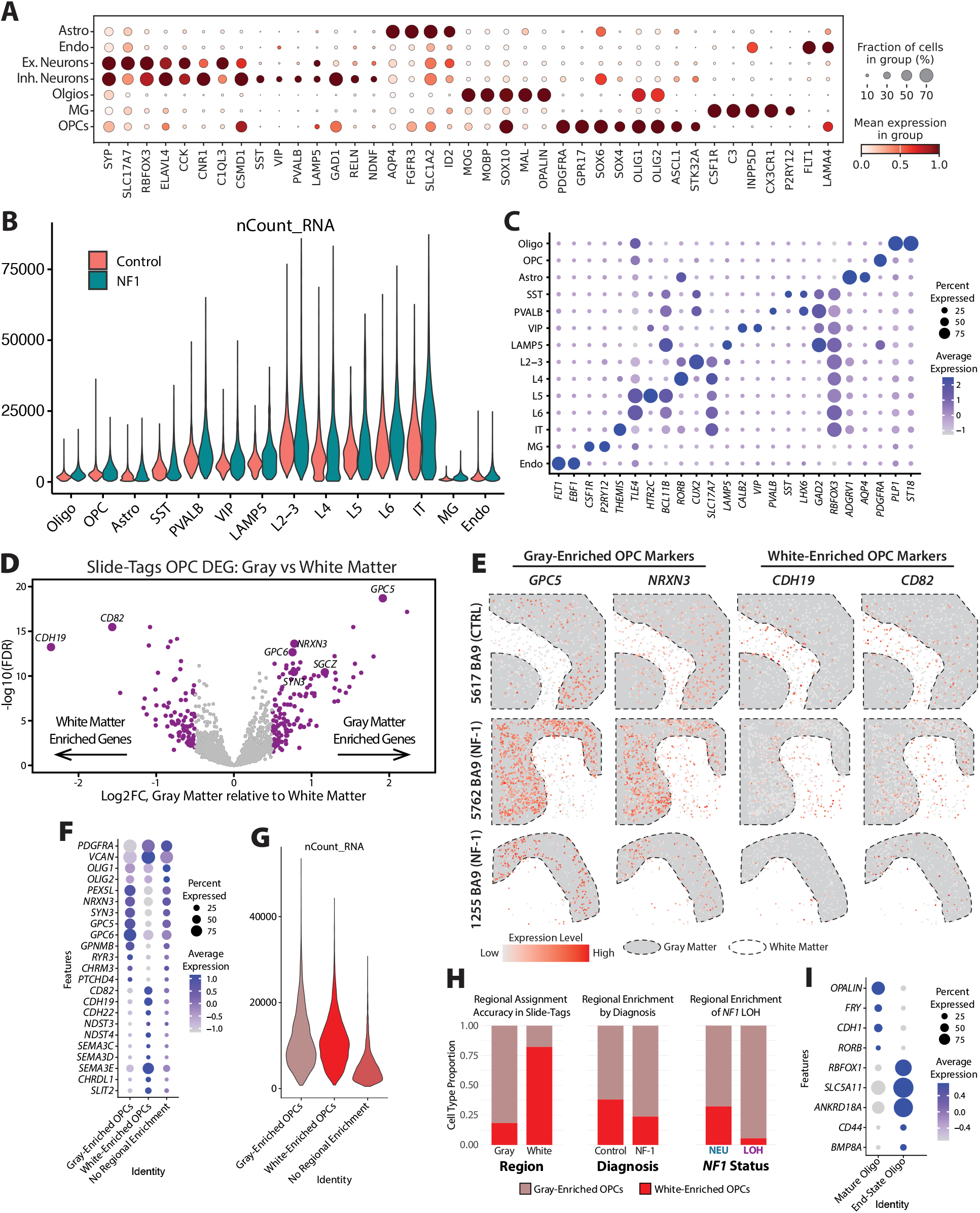
Spatial transcriptome datasets sequencing and clustering quality metrics. (**A**) Dot plot showing marker gene expression in all cell types in MERFISH dataset. (**B**) Violin plot of UMI count across cell types in Slide-Tags dataset, split by NF-1 diagnosis. (**C**) Dot plot showing marker gene expression in all cell types in Slide-Tags dataset. (**D**) Volcano plot of region-specific DEGs in OPCs from spatial snRNA-seq data. (**E**) Expression of representative gray/white matter-enriched OPC marker genes. (**F**) Dot plot showing expression of region-specific OPC marker genes across subtypes. (**G**) Violin plot of nCount RNA across gray/white-enriched OPCs showing that the cells with no clear regional enrichment have lower number of transcripts captured. (**H**) Stacked bar plots showing the proportion of transcriptionally determined gray/white-enriched OPCs in (left) gray and white matter in spatial snRNA-seq data, (middle) control and NF-1 donors, and (right) cells with chr17 LOH or neutral. (**I**) Expression of marker genes for OL subtypes.

**Figure S6:**
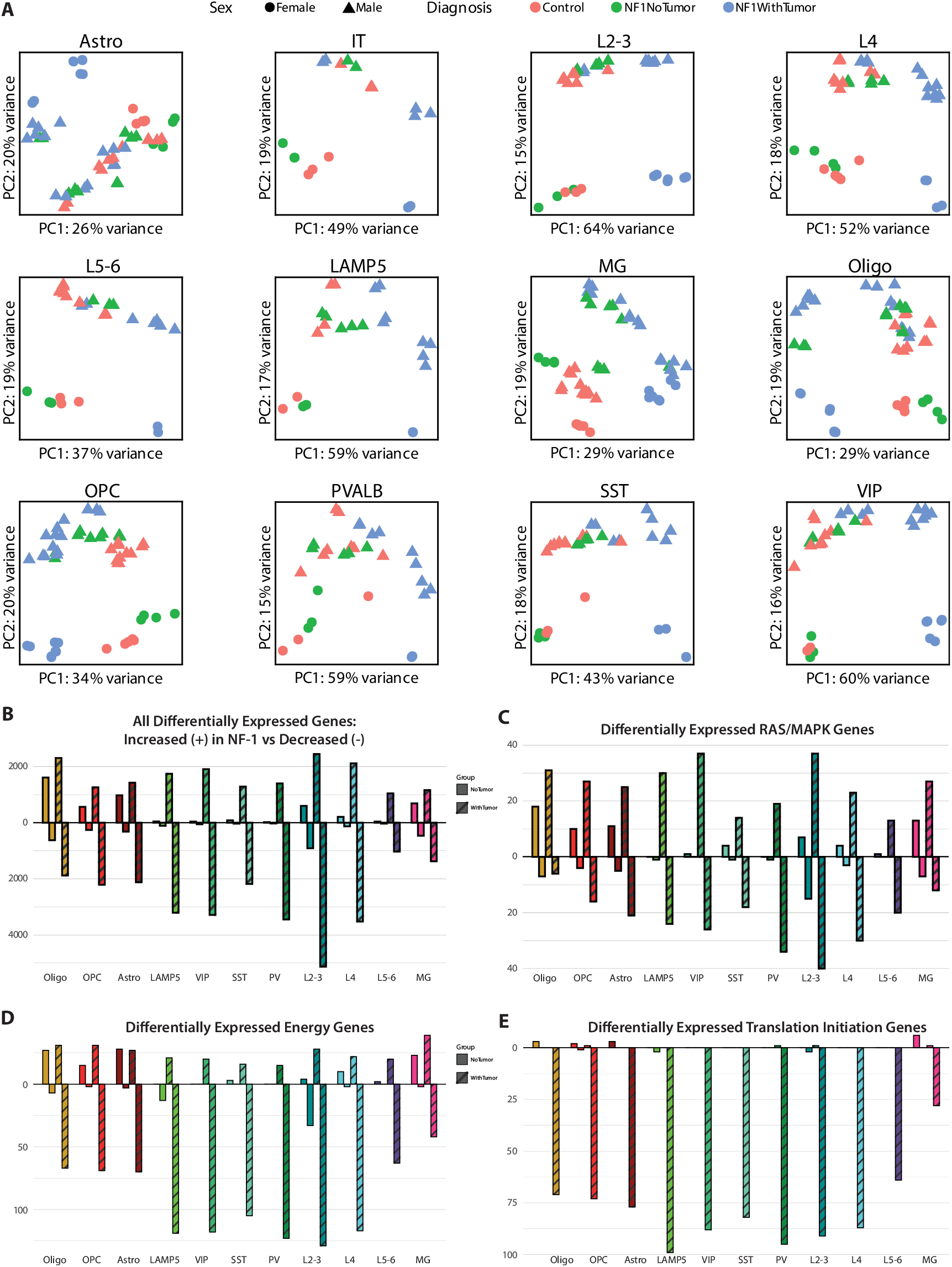
Principal component clustering of NF-1 donors harboring a brain tumor and corresponding transcriptional differences. (**A**) Principal component analysis shows all samples across all cell types cluster by sex as expected, but additionally, NF-1 samples from individuals with brain tumors cluster in distinct patterns. PCA generated from DESeq2 normalized counts from each cell type group. Cell Type Abbreviations: Oligo=Oligodendrocytes, OPC=Oligodendrocyte Progenitor Cells, Astro = Astrocytes, LAMP5 = Lysosomal Associated Membrane Protein Family Member 5 Interneurons, VIP= Vasoactive Intestinal Peptide Interneurons, SST = Somatostatin Interneurons, PV = Parvalbumin Basket Cells, L2-3 = Cortical Layer 2/3 Excitatory Neurons, L4 = Cortical Layer 4 Excitatory Neurons, L5-6 = Cortical Layer 5/6 Excitatory Neurons, IT = Intratelencephalic Excitatory Neurons, MG = Microglia. (**B**) Number of differentially expressed (FDR < 0.05 & |log_2_(FC) | > 0.5) genes, as well as gene sets from the Reactome database plotted including (**C**) RAS/MAPK, (**D**) energy (glycolysis and oxidative phosphorylation) genes, and (**E**) translation initiation genes, revealing that NF-1 brain samples from individuals with brain tumors show multiple distinct gene expression programs.

**Figure S7:**
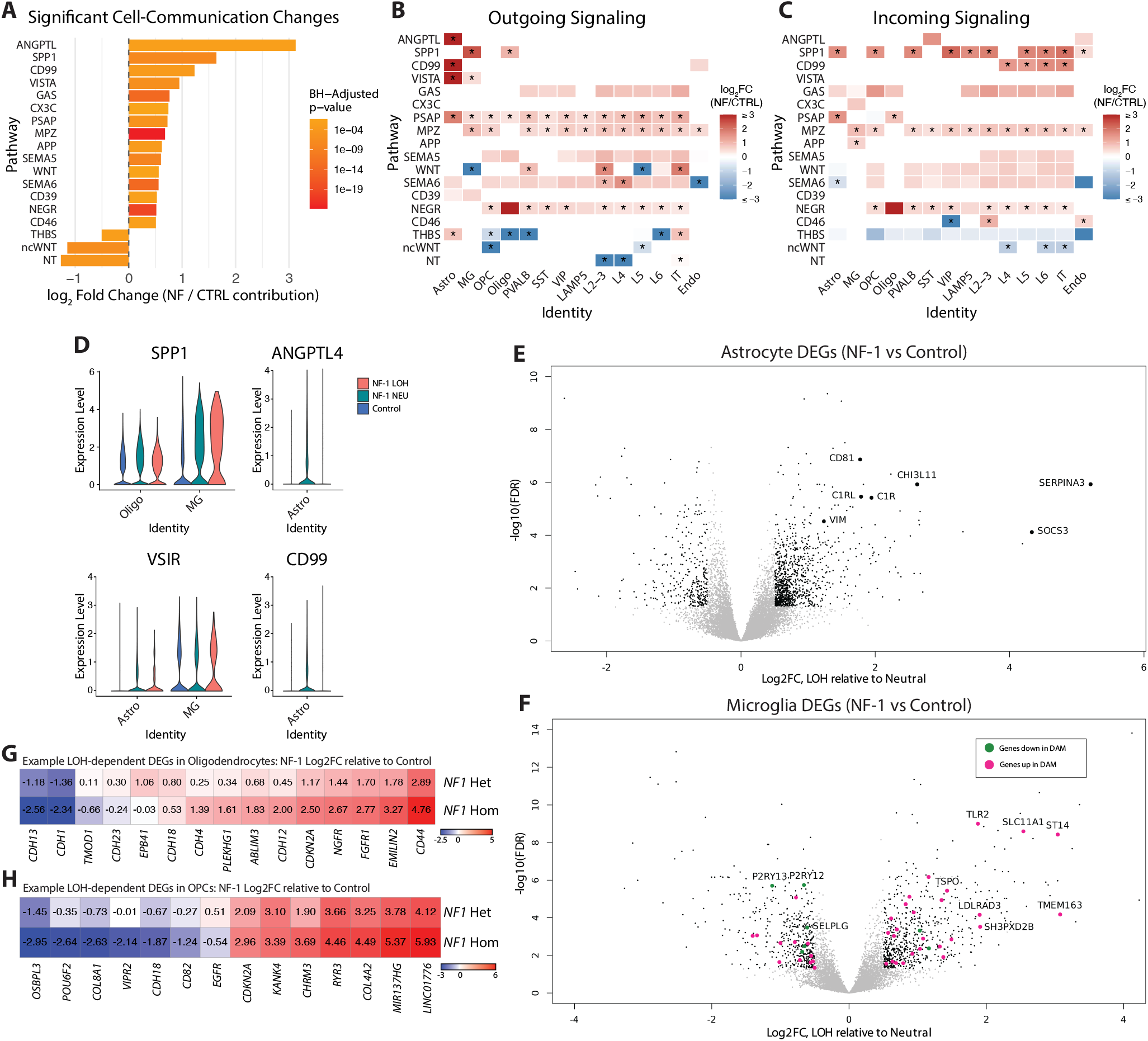
Cell-cell communication and DEG analysis highlight increased astrocyte reactivity and microglia activation in NF-1. (**A**) Pathway-level enrichment showing overall shifts in cell-cell communication in NF-1 donors vs controls. (**B**) Heatmap showing pathway enrichment for outgoing signaling (ligand-producing cells) and (**C**) incoming signaling (receptor-producing cells) (* p < 0.05 by random permutation testing, Benjamin-Hochberg adjusted). (**D**) Violin plots showing ligand receptor expression from the top four differential pathways in A. (**E**) Volcano plots displaying DGE results in NF-1 without brain tumor astrocytes vs neurotypical OLs, revealing increased reactive astrocyte genes in NF-1. (**F**) Volcano plots displaying DGE results in NF-1 without brain tumor microglia vs neurotypical microglia, revealing increased Disease Associated Microglia (DAM)-like gene expression patterns in NF-1, DAM genes from Keren-Shaul et al. (**G**) Heatmap of significant DEGs in LOH vs Neutral OL DGE demonstrating greater changes in *NF1* LOH OLs than *NF1* Neutral OLs relative to neurotypical OLs. (**H**) Heatmap of significant DEGs in LOH vs Neutral OPC DGE demonstrating greater changes in *NF1* LOH OPCs than *NF1* Neutral OPCs relative to neurotypical OPCs.

**Figure S8:**
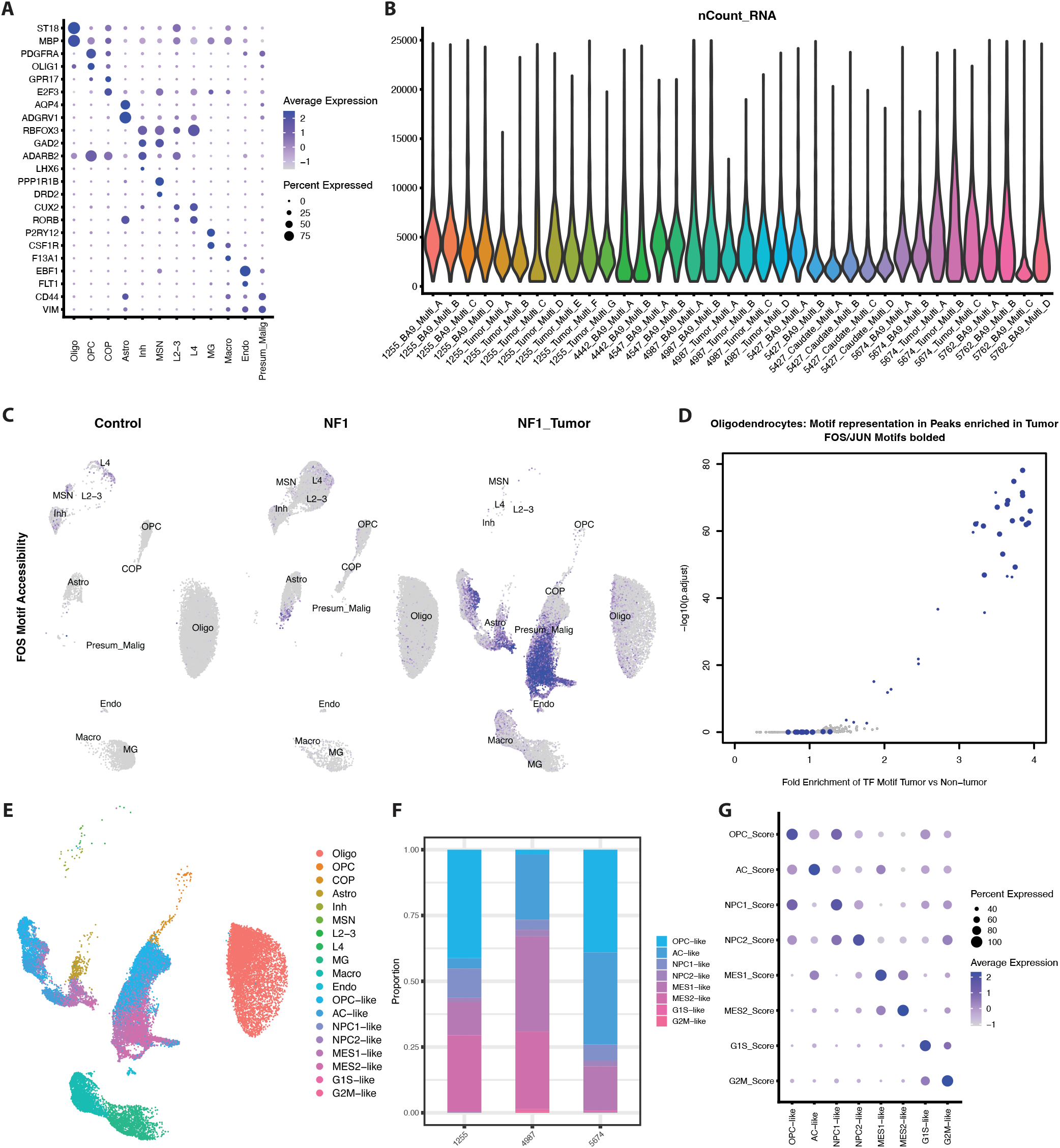
Marker gene expression, cell state distribution, UMI count, and FOS/JUN motif accessibility in multiome dataset. (**A**) Dot plot showing marker gene expression in all cell types in multiome dataset. Cell Type Abbreviations: Oligo=Oligodendrocytes, OPC=Oligodendrocyte Progenitor Cells, COP = Committed Oligodendrocyte Progenitors, Astro = Astrocytes, Inh = Inhibitory Neurons, MSN = Medium Spiny Neurons, L2-3 = Cortical Layer 2/3 Excitatory Neurons, L4 = Cortical Layer 4 Excitatory Neurons, MG = Microglia, Macro = Macrophages, Endo = Endothelial Cells, Presum Malig = Presumed Malignant Cells. (**B**) Violin plot of UMI count across cell types in multiome dataset. (**C**) Feature plot split by sample type showing FOS Motif accessibility greatly increased in presumed malignant nuclei but subtly increased in tumor OLs. (**D**) Enrichment of transcription factor motifs in ATAC peaks more accessible in tumor samples than matched non-tumor samples in OLs; FOS and JUN motifs strongly enriched in tumor-enriched ATAC peaks in OLs. (**E**) UMAP representation of tumor samples, including cell state for presumed malignant nuclei. (**F**) Stacked bar chart showing the distribution of cell states in presumed malignant nuclei across the 3 tumors. (**G**) Dot plot showing expression of cell state modules across malignant cell states.

**Figure S9:**
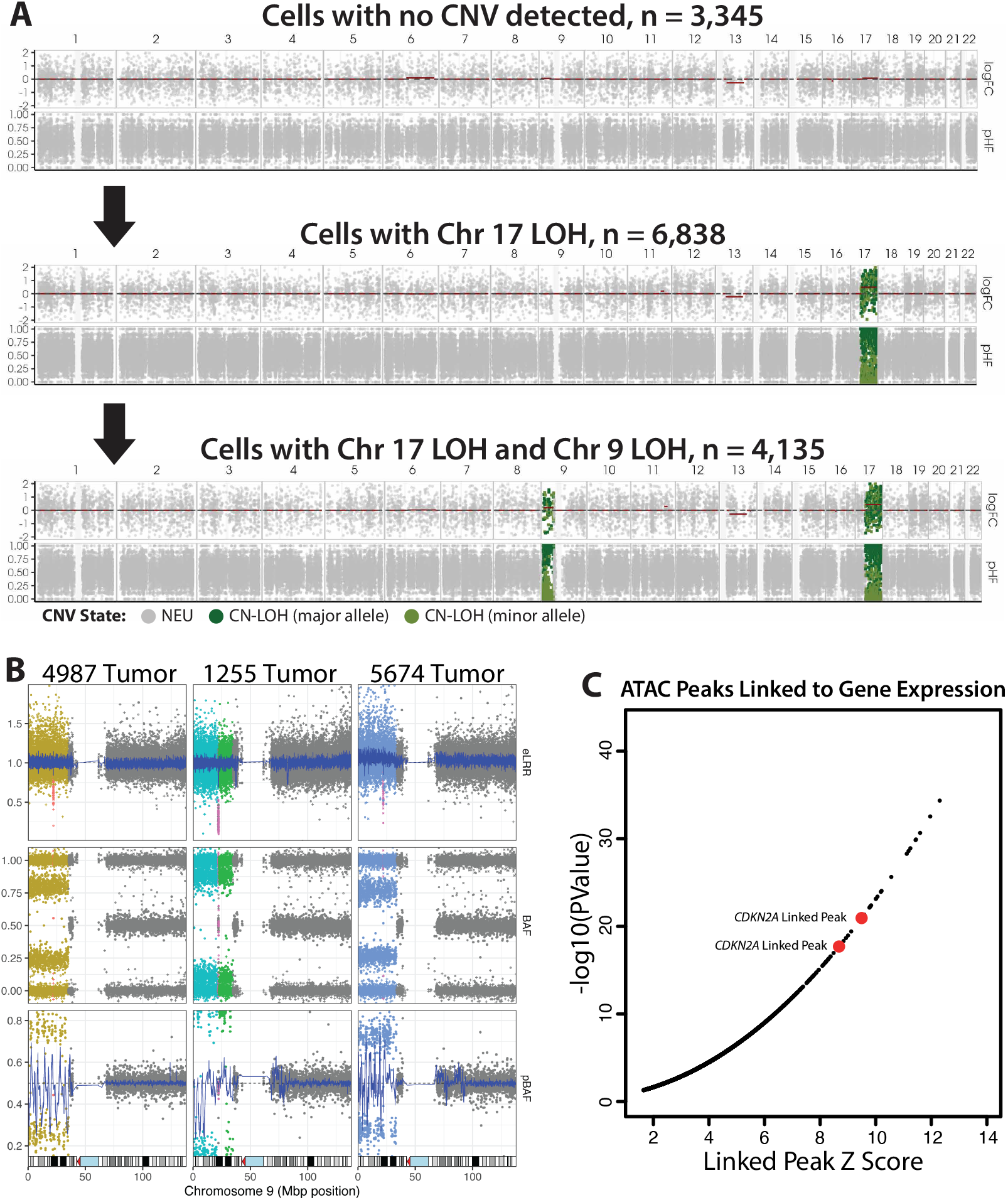
*CDKN2A* loss occurs after Chromosome 17 LOH and leads to chromosome 9 loss of heterozygosity in all 3 NF-1 tumors. (**A**) Pseudobulk of snRNA-seq data for three clones identified by Numbat in the tumor from donor 1255. No mCA were identified in the top clone. The middle clone has a single mCA on chromosome 17 which is also present in the bottom clone in addition to an mCA on chromosome 9 overlapping *CDKN2A*. (**B**) MoChA plots showing *CDKN2A* loss in all three NF-associated tumors. The top 3 plots show genotyping intensity and show a drop in intensity corresponding to the location of *CDKN2A* indicating a deletion. In addition to the focal deletion in *CDKN2A* there is also a CN-LOH event encompassing the majority of chr9p including *CDKN2A* which drives the deletion to homozygosity. (**C**) All chromatin peaks significantly linked (p < 0.05) to changes in gene expression, with two peaks in *CDKN2A* gene in the top 30.

## Materials and Methods

### Human Tissue Sources

Post-mortem frozen human tissues were obtained from the NIH Neurobiobank according to approved institutional protocols (table S1). All samples were shipped on dry ice and were stored at −80°C prior to use.

### Hematoxylin and eosin staining

Fresh-frozen brain sections (10 µm) were fixed in 80% acetone for 1 minute. Sections were stained with Gill 3 hematoxylin for 1 minute, rinsed in water for 2 minutes, immersed in ammonia water for approximately 5 seconds, and rinsed again in water for 2 minutes. Sections were subsequently stained with 1% alcoholic eosin for 3 minutes, dehydrated in 95% ethanol for 1 minute and in two changes of 100% ethanol for 1 minute each, cleared in two changes of xylene for 2 minutes each, and coverslipped with Permount mounting medium.

### Ki-67 (MIB-1) Staining

The IHC staining was performed on previously unfixed frozen human brain tissue. Slides were cut and held at −80°C until ready to be stained. The unfixed frozen brain slides were left to air dry at room temperature for 20 minutes to kick off the staining process. After air drying, the sections were placed into 4% PFA for 20 mins at room temperature for fixation. This was followed by 3 washes in TBST (TBS buffer with Tween 20 detergent) and then 1 wash in TBS (all washes are 5 mins each for the whole process). After washing, tissue sections were permeabilized using 3% BSA + 0.05% TX-100. The slides were then put through another sequence of washes – TBS, TBST, then TBS.

Sections were incubated with 3% BSA for 1 hour at room temp. Sections were incubated with Ki-67 (MIB-1), Cell Signaling Tech, Cat. #: CST 12202, @ 1:500) overnight at 4°C. The slides were washed with TBS, TBST, and TBS before incubating with goat anti-rabbit IgG HRP secondary antibody (Abcam, Cat. #: ab214880, 1:2 in 2% BSA) for 90 minutes at room temperature. Samples were then washed again with TBS, TBST, and TBS. The slides were developed in DAB (Diaminobenzidine) peroxidase substrate kit (Vector Labs) and counter-stained with hematoxylin. After dehydration, the slides were mounted with Permount.

### Evaluation for evidence of neoplasms

A board-certified neuropathologist evaluated H&E and MIB-1 stains for features of neoplastic infiltration. Tissue was considered free of overt tumor involvement when sections lacked morphologic hallmarks of infiltrating glioma (including hypercellularity, nuclear atypia, mitotic figures, microvascular proliferation, and necrosis) and showed low MIB-1 labeling restricted to rare nuclei consistent with non-neoplastic brain.

### Bulk DNA Extraction

Bulk DNA extraction was performed on fresh-frozen post-mortem human brain tissue using the DNeasy Blood & Tissue Kit (Qiagen catalog# 69504) as per the manufacturer’s instructions. 250mg of tissue, previously stored at −80°C, was scraped on dry ice into a 1.5 mL Eppendorf Lo-Bind tube with Buffer ATL and proteinase K. The protocol was then continued as per the manufacturer’s instructions and yield was quantified using the Qubit 1X dsDNA, high sensitivity assay kit (ThermoFisher catalog# Q33231).

### Bulk Short-Read Sequencing

A total of 200-500ng of bulk extracted DNA was used for PCR-free library prep with the Kapa HyperPrep Kit (Roche catalog# 07962371001) and Kapa Unique Dual-Indexed Adapter Kit (Roche catalog# 08861919702). Prepared libraries were sequenced on either Illumina NovaSeq 6000 or NovaSeq X Plus targeting mean coverage of 30x across the genome. Read preprocessing, alignment, and germline variant calling were performed using nf-core/sarek v3.1.1^81^ using Haplotype Caller ^82^ for germline variant calling and SnpEff ^83^ for variant annotation.

### Bulk Long-Read Sequencing

High molecular weight (HMW) DNA was extracted from 25mg of brain tissue with the Nanobind PanDNA kit (PacBio catalog# 103-260-000) following manufacturer’s instructions. Extracted HMW DNA quality was assessed with the Femto Pulse system using the Genomic DNA 165kb Kit (Agilent catalog# FP-1002-0275) to confirm the presence of a strong peak at 150kb. A total of 2ug of high quality HMW DNA was used for PacBio library preparation using the SMRTbell prep kit 3.0 (PacBio catalog# 102-141-700). Libraries were sequenced on a PacBio Revio system with a single sample per SMRT cell yielding 30-40x coverage across the genome with HiFi read length N50 > 15kb. Reads were aligned to GRCh38 using the pbmm2 wrapper of minimap2 using the HiFi preset.

### Variant Phasing

To achieve chromosome-scale phase information we used a multi-step approach combining statistical and read-backed phasing followed by correction of phase-switch errors on chr17q enabled by samples with high-frequency LOH on chromosome 17q. First, we performed statistical phasing on each sample using SHAPEIT4 v4.2.2^84^ with default settings inputting germline variants as called from bulk short-read sequencing and using the 1000 genomes project dataset (release 20190312) as the population reference ^85^. This provides chromosome-scale phase; however, this method is restricted to common variation present in the population reference which precludes establishing the phase of the germline pathogenic variants in *NF1*. To achieve chromosome-scale phase including the pathogenic *NF1* variants, we input long-read sequencing data to WhatsHap phase v2.0^86^ using default settings with an additional input of the SHAPEIT4 phased VCF. Using this strategy WhatsHap leverages the long-read data to assemble small local haplotypes, then finds variants within the local haplotype that overlap with the input phased VCF allowing the local haplotype to be scaffolded onto the chromosome-scale haplotypes from SHAPEIT4.

The above strategy yields a chromosome-scale phased VCF that includes the pathogenic germline *NF1* variants; however, the use of statistical phasing inevitably results in phase-switch errors ^13^ which reduce the sensitivity of our downstream analyses. To partially resolve phase-switch errors we selected samples harboring mCA with high frequency BAF deviations on chromosome 17 detected by MoChA (version 2024-09-27) ^13,14^. To determine the breakpoints of phase switch errors, we input phased BAF values from MoChA that fall within the mCA region into a Hidden Markov Model (HMM) using the R package depmixS4 v1.5-1^87^. We fit the data to a two-state HMM run with default settings and selected the model that maximized log likelihood across 10 iterations. Regions between transition points were converted to a BED file and regions containing less than 5 heterozygous markers were removed from the final set. The resulting BED file was used to reverse the phase of variants inside every other phase block using bcftools v1.22^88^ to orient the phase relative to the allele harboring the pathogenic *NF1* variant.

### Bulk DNA mCA Detection

To identify mCA we submitted 200ng of extracted DNA per sample to Psomagen (Rockville, MD, USA) to be run on the Infinium Global Screening Array v3 (Illumina catalog# 20030770). Resulting IDAT files were converted to GTC format using bcftools plugin idat2gtc (version 2024-09-27) and subsequently converted to BCF format on reference GRCh38 using plugin gtc2vcf (version 2024-09-27) ^89^. We generated an initial mCA call set by phasing BCF files exclusively with SHAPEIT4 using 1000 genomes project data as a reference after which the phased BCF was input to MoChA run with default settings and recommended call filtering. The initial call set was used to identify mCA on chromosome 17q with high BAF deviations which were used to correct phase switch errors (see Variant Phasing). The final mCA call set was generated using the phase information derived from the combination of statistical phasing, read-backed phasing, and phase switch correction.

### Bulk DNA Point Mutation Detection

DNA sequencing libraries were prepared using 100ng of bulk extracted DNA input to the SureSelect XT HS2 library preparation kit (Agilent catalog# G9983A) with molecular barcode (MBC) tagged adaptors that differentially tag the Watson and Crick strands of DNA enabling duplex consensus calling which improves variant calling sensitivity and specificity. A total of 500-1000ng of prepared libraries were input into hybridization-based capture reactions (one per sample) using a custom panel targeting 201 genes including *NF1* (Agilent, table S2). Final target-capture libraries were sequenced on a NovaSeq X Plus to a mean depth of 21594x resulting in a mean depth of 416x after duplex consensus was performed.

Data preprocessing was performed using the Agilent Genomics ToolKit (AGeNT v3.1.2) and the Consensus Read Kit (CReaK v3.1.2). Briefly, the trim command in AGeNT was used to trim sequencing adaptors from reads and add an annotation with the MBC sequences from each read pair. Following trimming the reads were mapped to GRCh38 with bwa mem v0.7.18-r1243-dirty. After mapping the reads, CReaK was used to group reads originating from the same molecule of DNA and identify the strand of origin based on both alignment position and MBC information. CReaK was run in both duplex (exclusively retain consensus reads from molecules with both strands) and hybrid (retain all consensus reads) modes. Output BAM files for both duplex and hybrid consensus modes were input for variant calling using nf-core/sarek v3.1.1 running Mutect2^90^ and Variant Effect Pressdictor (VEP) ^91^ with the `--joint mutect2` and `--wes` settings as well as the `--intervals` setting to limit analysis to regions included the targeted panel. For Mutect2 paired tumor/normal analysis the bulk short-read sequencing data from each individual was used as the normal. Variants were filtered to include only those listed as PASS which VEP predicted to have moderate or high impact on *NF1*.

### Nuclei Isolation from Human Brain Tissue

25mg of fresh-frozen postmortem human brain tissue, previously stored at −80°C, was scraped on dry ice into a 7 mL Dounce homogenizer on ice with 1 mL of chilled nuclear isolation media (250mM Sucrose, 25mM KCl, 5mM MgCl_2_, 10mM Tris-HCl pH 8, 0.1% Triton X-100, 0.2U/ul RNase inhibitor, 1 tablet per 10mL of protease inhibitor, and 1mM DTT). Nuclei were then pelleted after 40 µm filtration and centrifugation at 900G for 10 mins or pelleted after layering of homogenate onto a sucrose gradient with centrifugation at 30,000G for 1 hr. Pellets were then resuspended in 500 µL cold blocking buffer (1X PBS supplemented with 0.8% Bovine Serum Albumin, 0.2U/ul RNase inhibitor, 1 tablet per 10mL of protease inhibitor, and 1mM DTT) for 10 mins on ice. Nuclei were then stained accordingly with the following conjugated antibodies: anti-NeuN Alexa Fluor 488 (Sigma Aldrich catalog# MAB377X) at 1:500, anti-Sox10 DyLight 550 (Novus Biologicals catalog# NBP2-59621R) at 1:500, DAPI (ThermoFisher Scientific catalog #62248) at 3:500. Nuclei staining with the conjugated antibodies occurred for 30 mins at 4°C with protection from light. The nuclei pellet was then centrifuged at 900G for 5 mins, supernatant discarded, and the pellet was washed in 500 µL cold blocking buffer 1-2x. Pellet was then centrifuged at 400G for 5 mins, supernatant discarded, and the remaining pellet was resuspended in 500µL cold blocking buffer, and filtered through 40 µm filter into a fresh, cold 1.5 mL Eppendorf Lo-Bind tubes for sorting.

### Fluorescence Activated Nuclei Sorting

Fluorescence activated nuclei sorting using flow cytometry was performed on DAPI positive nuclei, and in select cases, gated for neurons and oligodendroglial lineage cells. DAPI positive 2N nuclei were targeted for sorting with exclusion of doublets using forward and side scatter-height and width gating. Neurons were sorted by targeting large NeuN positive nuclei with the highest forward Scatter Area (FSC-A) ^92^ or more broadly by targeting NeuN+/Sox10-population. Oligodendroglial lineage cells were largely sorted by targeting the Sox10+/NeuN-population.

### Single-Nucleus DNA Sequencing

To identify both somatic point mutations and somatic mCA in single cells, we used targeted single-nucleus DNA sequencing (scDNA-seq) using the Tapestri system (MissionBio) with the DNA v3 chemistry. Following nuclei isolation from post-mortem human brain tissue nuclei were stained with DAPI and used for fluorescence activated nuclei sorting to collect 200,000-300,000 DAPI+ nuclei. Nuclei were centrifuged at 4°C and 500G for 10min and the resulting pellet was isolated and resuspended in 36µL of Cell Buffer (MissionBio). The concentration of nuclei was determined by counting DAPI+ nuclei with a hemocytometer on an EVOS M5000 (Thermo Fisher) to ensure the frequency of nuclei aggregates was <5% and the concentration of nuclei was >3000 nuclei/µL. Library preparation was performed according to the manufacturer’s protocol using a custom primer panel targeting the exons of *NF1* in addition to 262 common polymorphic sites across chromosome 17 with increased marker density across the gene body of *NF1* and 2Mb of flanking sequence (table S5). Resulting libraries were sequenced on a NovaSeq X Plus targeting 80x reads per amplicon per cell.

Data preprocessing was performed using the Tapestri analysis pipeline v3.4 (MissionBio) with default settings yielding cell x variant matrix. Variant filtering and chromosome 17 LOH calling was performed using the Mosaic v3.12.2 (MissionBio). We applied both cell-level and variant-level filters to remove low quality data using provided filtering functions in Mosaic with default settings (`filter_variants`, `filter_variants _consecutive`, and `filter_barcodes`).

Point mutations were filtered to remove germline variants present in the paired bulk short-read sequencing data and Mosaic filter `filter_somatic variants` with the option `min_mut_prct_cells_denovo=0.5` to remove calls present in less than 0.5% of cells in the sample. In addition to default Mosaic filters, we applied a cross-sample filter to remove point mutations that were present in >0.5% of cells in multiple samples. For tracking clonal structure of somatic variants impacting *NF1* we subset these variants to those VEP predicted to have moderate or high impact on *NF1*.

To identify LOH events impacting chromosome 17 in individual cells we started with the set of variants surviving default Mosaic filters described above (not including `filter_somatic_variants`) and subset to variants on chromosome 17 that were heterozygous in >30% of cells within the sample. The resulting cell x variant matrix was input to Mosaic `find_clones` with `similarity = 0.95` to identify clones of cells where germline heterozygous variants are instead homozygous (LOH). LOH breakpoints were determined for each cluster and per-cell LOH calls were made based on cluster membership.

### Single-Nucleus RNA Sequencing

Nuclei were sorted into 1.5 mL Lo-Bind tubes containing 10x GEM master mix supplemented with nuclease-free water, as described in the 10X Genomics Chromium Single Cell 3’ Reagent Kits v3.1 User Guide (CG000315 Rev E). GEM generation & barcoding, post GEM-RT cleanup & cDNA amplification, and 3’ gene expression library construction was largely performed per the manufacturer’s instructions and quality control measurements on the libraries were performed using the Agilent 4200 TapeStation System prior to sequencing. Paired-end sequencing of the libraries was then performed on Illumina NovaseqX platform by Psomagen or Novogene targeting 50,000 reads per nuclei.

### Single-Nucleus RNA Preprocessing

Gene counts matrices for our data were generated by aligning the raw FASTQ files to the GRCh38 reference human genome using Cell Ranger v.9.0.1 (10X Genomics) and the reference file “gex-GRCh38-2024-A” with default settings, including intronic reads. Seurat v.5.3.0^93^ R package was used for all subsequent downstream analyses. Libraries from experiments using different sort strategies (i.e. *SOX10* or *NEUN* staining) were processed separately. Standard QC was performed including 1] percent mitochondrial gene expression <5%, 2] >200 but <10,000 unique genes (nFeature RNA) to exclude doublets and low quality nuclei. High quality nuclei data were then normalized (NormalizeData) and scaled (ScaleData) according to default Seurat processing parameters. CCA Integration was then performed based on donor identity to avoid donor-specific or dataset-specific clustering. Dimensionality reduction was performed using PCA (RunPCA), and the top 17 principal components were used for downstream analysis based on visual inspection of an elbow plot of the proportion of variance captured by each PC for the unsorted snRNA-seq experiments. For *SOX10* sorted experiments, decreased cell type complexity meant that 12 principal components were necessary. For *NEUN* sorted experiments and the spatial snRNA-seq dataset, 18 principal components were used. For OPC and OL sub-clustering with all datasets combined, 15 principal components were used. Upon clustering, more doublets were removed based on expression of marker genes from multiple cell types in the same nuclei. UMAP (RunUMAP) was used for dimensionality reduction and visualization. Annotation was performed manually based on known cluster markers used in published human brain datasets ^25,94^.

### Single-Nucleus RNA mCA Detection

To detect mCA in single cells using snRNA-seq, we applied Numbat v1.4.2^16^ which uses information from both expression level and phased allele frequency shifts to genotype single cells for known CNVs. Numbat was run using a custom expression reference created using snRNA-seq data from DAPI+ nuclei sorted from control samples. We also used the `segs_consensus_fix` option in Numbat to input the high confidence mCA calls for each sample obtained from microarray data and decreased the SNP population allele frequency from >5% to >0.5%. Otherwise, Numbat was run with default settings. The mCA status for each cell was determined from the `joint_posts.tsv` output from Numbat containing the posterior probability that each cell harbors each mCA or that no mCA are present. An mCA was called as present if the posterior probability for the given cell (`p cnv`) was ≥ 0.99. Similarly, a cell was only called as neutral if the posterior probability (`p neu`) was ≥ 0.99. Cells where `p cnv` and `p neu` were <0.99 were listed ‘no call’. To estimate the false-positive rate for this approach, we ran Numbat on control samples using a `segs_consensus_fix` listing chromosome 17q copy-neutral LOH. These results revealed cell type specific variation in false-positive rate which was incorporated into our analysis of chromosome 17 mCA burden in NF samples. In burden quantification, only cell type groups within individual samples that had >20 nuclei were included to minimize the noise inherent in burdens in a small sample size.

To determine the haplotype lost from LOH events in NF-1 samples, we used donor-specific phased VCFs (see Variant Phasing) to derive haplotype-specific read counts for input to Numbat. First, phased VCFs were subset to chr17 and cellsnp-lite (v1.2.3) was run on the snRNA-seq BAM files to provide cell-level read counts for all phased heterozygous sites on chr17. The cellsnp-lite pileup output was merged with the original phased VCF to build a Numbat-compatible allele-count table which was input to Numbat as described above. For each phased heterozygous SNP, reference and alternate read counts were mapped to haplotype 1 or haplotype 2 according to the phased genotype. Note the phased VCF files for each NF-1 donor are oriented such that haplotype 1 contains the pathogenic germline *NF1* variant. For each cell, haplotype 1 and haplotype 2 reads (UMI-deduplicated) were summed across all informative SNPs in the segment-fix region. Cells with greater haplotype 1 total reads were labeled haplotype 1, cells with greater haplotype 2 total reads were labeled haplotype 2, and cells with equal or missing informative support were labeled undetermined.

### Differential Gene Expression Analysis

EdgeR v4.4.2^95^ with default parameters was used on sample-bulked raw counts data for each cell type for all differential gene expression analysis, with age and sex effects regressed out. Benjamini-Hochberg correction was applied to generate FDR values on all nominal p values for genes, and genes were considered significantly changed if the FDR was less than 0.05 and the absolute value of the log_2_FC was greater than 0.5. For LOH vs neutral differential gene expression, a paired differential gene expression analysis was performed for LOH and neutral nuclei within each sample and cell type group. All NF-1 samples with LOH detected were included for this analysis. For all analyses, as in the LOH burden calculation, only cell type groups within a sample with >20 nuclei were used. For Principal Component Analysis (PCA), DESeq2 v1.46^96^ was used with default parameters to normalize counts and extract principal components.

### Cell-Cell Communication Analysis

We employed the CellChat v2.2.0.9001^29^ to evaluate cell-cell communication networks using the extensive database of known human ligand-receptor interactions as defined in CellChat v1 to remain consistent with prior analyses. We ran CellChat with default parameters for dataset comparison separately running cells from control and NF-1 donors prior to merging objects for subsequent comparison. CellChat applies the law of mass action to the mean expression level of paired ligands and receptors to identify cell communication pathways that differ between NF-1 and control donors. The significance of interactions is determined by random permutation statistical testing and subsequently adjusted with Benjamin-Hochberg correction to control for multiple hypothesis testing. We required that ligand-receptor pairs be identified as significant in at least 3 pairs of source-target cell types and focused our analysis on pathways with adjusted p-values above 0.05, log_2_FC above 0.5, and total signaling contribution above 0.1.

### MERFISH Sample Preparation and Imaging

Fresh-frozen post-mortem human brain tissue was embedded in OCT and then sliced at 10 µm thickness in a cryostat. Slices were then placed directly onto Vizgen MERFISH slides, placed in a 21.2cm^2^ tissue culture dish, and incubated at −20°C inside the cryostat for 15 mins. Slides were then transferred to room temperature for 15 mins to allow the tissue to better adhere to the slide. Slides were fixed for 15 mins using 4% Paraformaldehyde in PBS and permeabilized overnight using 70% ethanol. Autofluorescence quenching was performed the next day, and the probe hybridization, gel embedding, and clearing were largely performed as described in the MERSCOPE Fresh and Fixed Frozen Tissue Sample Preparation User Guide (91600002 Rev F) with a few modifications including changing clearing solution on day 4 or 5 after initial addition, and imaging up to about 9 days after gel embedding as long as the integrity of the gel was stable enough for imaging. Samples were stained with DAPI and PolyT and imaging performed as per the MERSCOPE Instrument User Guide (91600001 Rev G).

### MERFISH Analysis

Cell segmentation was carried out using automated Watershed option as noted in the MERSCOPE Instrument User Guide (91600001 Rev G). MERFISH count matrices and metadata were processed on a per-sample basis using Scanpy (v.1.10.4). Cells were filtered by removing cells with total counts below the sample specific 10th percentile and cells with fewer than 10 expressed genes, after which data were normalized, log-transformed and scaled. For sample integration, PCA was performed and batch effects were corrected using Harmony, and neighbor distance and UMAP embeddings were constructed from the Harmony corrected space, followed by Leiden clustering at resolution 0.15. Cluster marker genes were identified using Wilcoxon rank-sum tests and used to annotate clusters into major brain cell types, with annotations further supported and validated from a published MERFISH reference ^97^. To categorize white and gray matter regions, the integrated dataset was converted to a Seurat object (v.4.3), and module scores were computed using representative white matter (MOG, OLIG1) and gray matter (SLC17A7, GAD1) markers with the AddModuleScore function. Cells were assigned to the higher scoring category first and then spatially smoothed using a k-nearest-neighbor (KNN) majority vote based on the spatial coordinates (k=750). Differences in cell type composition between NF-1 and control samples were assessed separately within white and gray matter regions using scProportionTest in scDown ^98^, with cell types as cell groups and sample conditions, i.e., NF-1 and control, as the comparison variables.

### Slide-tags Sample Preparation and Sequencing

Slide-tags was performed using the commercial Trekker Single-Cell Spatial Mapping Kit (Takara catalog# SK017) according to manufacturer protocol, with addition of a myelin removal step immediately following tissue dissociation (Minute Single Nucleus Isolation Kit for Neuronal Tissues/Cells, Invent Biotechnologies, catalog# TBS8031). Briefly, samples were embedded in OCT and then slice at 30µm thickness in a cryostat. Slices were then placed directly onto a 1cm x 1cm Trekker tile for processing. Sections immediately before/after the section used for Trekker were also collected and used for H&E staining. Immediately following sectioning, UV cleavage buffer was immediately added to the tile, and the tile was treated with UV for 7.5min with the manufacturer-provided UV lamp (Takara catalog# K011). Following UV treatment, the tissue adhered to the tile was dissociated by trituration and nuclei were purified using the above-mentioned myelin removal kit. Purified nuclei were stained with DAPI and counted with a hemocytometer on a fluorescent microscope. Captured nuclei were split across 1-3 snRNA-seq reactions with up to 10,000 nuclei each. Our dataset from 15 sections was split across 37 snRNA-seq reactions. snRNA-seq was performed according to the manufacturer protocol for the 10x NextGEM 3’ v3.1 chemistry with Feature Barcode technology for Cell Surface Protein (see 10x document CG000317 RevE). During this protocol the Trekker spatial barcode library is amplified in place of the Cell Surface Protein library following minor modifications to the 10x protocol as detailed in the Trekker manufacturer protocol. These modifications include use of the 10x TT index set, reducing the volume input to library amplification to 1µL, and reducing the number of PCR cycles to 7. The resulting libraries were sequenced on Illumina NovaSeq XP 25B lanes to 50,000 reads per cell for the gene expression library and 5000 reads per cell for the spatial barcode library.

### Slide-tags Analysis

The gene expression data was processed and cell types identified as described in Single-Nucleus RNA Preprocessing and mCAs were called with Numbat as described in Single-Cell RNA mCA Detection. Each spatial barcode library was processed with the trekker-v1.3.0 pipeline provided by Takara and libraries derived from the same tile were merged with curiotrekker merger-v1.1.0. This preprocessing retains only nuclei with a single confident spatial coordinate, removing cells with zero or multiple coordinates (likely due to multiplets) and yields a Seurat object pairing high-confidence nuclei with a single spatial coordinate. Spatial coordinates, gene expression data, and mCA calls were merged into a single Seurat object per donor that was used for downstream analysis.

To categorize regions of white and gray matter we employed a hybrid approach first using automated segmentation based on module scores from representative gene sets for white matter (*MBP, MOG, MOBP, PLP1, MAG, CLDN11, ERMN*) and gray matter (*SLC17A7, SYN1, SNAP25, RBFOX3, GAD1, GAD2, CAMK2A*). As with MERFISH, cells were assigned to the higher scoring category first then spatially smoothed using KNN (k=200). Due to the decreased cell density of Trekker Slide-tags compared to MERFISH, in a subset of samples this approach left small areas of white matter in gray matter and vice-versa. To further improve the gray/white matter boundaries we additionally used InteractiveSpatialPlot from Seurat to refine gray and white matter boundaries with the boundary present in an overlayed H&E section taken directly adjacent to the section used for Trekker. Differences in cell type composition between NF-1 and control samples were calculated separately using scProportionTest ^99^.

To determine whether LOH rate was associated with shifts in cell type proportions in gray and white matter, we first fit linear models (proportion ~ %LOH * region) for each cell type. NF-1 diagnosis was not included as a covariate due to its collinearity with LOH rate and likelihood ratio tests confirmed that NF-1 diagnosis added no explanatory power beyond LOH rate (p=0.51), while LOH rate explained significant variance beyond NF-1 diagnosis (p=0.005). From these models the per-region slopes, and p-values were estimated using emtrends (emmeans v1.11.1). LOH-associated fold changes were computed as the ratio of observed mean proportion to the proportion predicted at zero LOH rate.

### Single-Nucleus Multiome (ATAC/RNA) Sequencing

Nuclei were sorted into 1.5mL Lo-Bind tubes containing Lysis Dilution Buffer (10mM Tris-HCl pH 8, 10mM NaCl, 3mM MgCl_2_, 1% Bovine Serum Albumin, 1mM DTT, and 1U/µL RNase Inhibitor). The number of nuclei sorted at this stage was usually 5-fold more than the planned input nuclei given for the transposition to account for losses during lysis and detergent incubation. After sorting, 50µL of 1X lysis buffer (10mM Tris-HCl pH 8, 10mM NaCl, 3mM MgCl_2_, 1% Bovine Serum Albumin, 1mM DTT, 1U/µL RNase Inhibitor, 0.1% Tween 20, 0.1% Nonidet P40 Substitute, and 0.01% Digitonin) was added to each sample. This was incubated on ice for 2 minutes and then spun in a bucket centrifuge at 4C for 5 minutes at 500g. Without disrupting the pellet, all but 5µL of supernatant was removed and 1X Diluted Nuclei Buffer (1X Nuclei Buffer, 1mM DTT, and 1U/µL RNase Inhibitor) was added. This was spun down at 4°C for 5 mins at 500g, and all but 7µL of supernatant was removed. The remaining nuclei were then counted by taking 2µL of the sample solution and adding this to 8µL of 1X Diluted Nuclei Buffer and loaded into a hemocytometer. Using an input of 10000-15000 nuclei, the Transposition step of the 10x Genomics Next GEM Single Cell Multiome ATAC + Gene Expression (CG000338 Rev A) was performed as described in the protocol. All subsequent steps were carried out as described in this user guide. After generating ATAC and RNA libraries, paired-end sequencing was performed on Illumina NovaseqX platform by Psomagen or Novogene targeting 50,000 reads per nuclei.

### Single-Nucleus Multiome Data Processing

Data filtering and cell type calling was performed using the same parameters as in the snRNA-seq datasets, with the following modifications. Instead of filtering by unique gene number (nFeature RNA), cells passing QC had to be between 500 and 25,000 unique molecular identifiers for RNA molecules (nCount RNA) and between 1,000 and 70,000 unique molecular identifiers for DNA molecules (nCount ATAC) in addition to having less than 5% mitochondrial genes. Clustering was performed using “weighted nearest neighbor” analysis to integrate both chromatin accessibility and transcriptome information for each nucleus. The native Seurat FindMultimodalNeighbors function with principal components of transcriptome information 1 through 9 and principal components of chromatin accessibility 2 through 30 were used (principal component 1 of chromatin accessibility is usually dependent on sequencing depth) When running FindClusters, the SLM (SNN-Louvain-Modularity) algorithm was used instead of the default Louvain algorithm, and resolution was set to 0.8 (the default). Subclustering of individual tumor presumed malignant nuclei was performed the same way but with principal components of transcriptome information 1 through 8 and principal components of chromatin accessibility 2 through 15. Nuclei were presumed malignant if the clusters they belonged to expressed malignant cell marker genes *CD44* and *VIM* and no other brain cell type marker genes. This was validated by checking for enrichment of non-chromosome 17 copy number variants (called by Numbat) in presumed malignant nuclei. To call glioblastoma cell states, module scores were calculated using the Seurat AddModuleScores function for each of the published cell states ^51^. Each nucleus was assigned a state based on the highest module score. Only nuclei called as presumed malignant from tumor samples were assigned cell states. The ATAC portion of the data was analyzed within the same Seurat object using R packages Signac v1.14.0^100^ and EnsDb.Hsapiens v86 v2.99.0. To analyze transcription factor motif accessibility, the JASPAR2020 database v0.99.10^101^ was used, as well as the TFBSTools R package v1.44.0^102^, with the tax group parameter set to “vertebrates”. To add motifs to the Seurat object, the Signac AddMotifs and RunChromVAR functions were used, along with the BSgenome.Hsapiens.UCSC.hg38 v1.4.5 database. Differentially accessible peak analysis was performed in the same manner as the differential gene expression, except the absolute value of the log_2_FC threshold was set to 1 instead of 0.5. The Signac LinkPeaks function with default parameters and the same genome database was used to identify significantly linked gene-peak combinations in the dataset.

For mCA calling from multiome data we applied Numbat on the RNA component of the dataset. Multiome analysis with Numbat used a custom reference built for multiome data using data from control donors. Similar to LOH calling from snRNA-seq we required a posterior probability of ≥ 0.9 to identify a cell as having a given mCA (from p cnv) or lacking the mCA (p neu). The mCAs identified from microarray data were used for genotyping using the `segs_consensus_fix` option in Numbat. Microarray data revealed chr9 CN-LOH secondary to, and overlapping with focal deletions of *CDKN2A* in all tumor samples. The multiome data was not sufficiently powered to detect the small focal deletions of *CDKN2A*, therefore the larger CN-LOH events encompassing the entirety of chr9p were used to identify cells with homozygous loss of *CDKN2A*.

